# SPG11 models reveal lysosomal calcium as a regulator of neural progenitor proliferation

**DOI:** 10.1101/2025.07.21.665525

**Authors:** Dominic Samaroo, Margaux Cauhape, Léa Bernachot, Frédéric Darios

**Author notes:** **Correspondance**: Frédéric Darios, Paris Brain Institute - ICM, Hôpital de la Salpêtrière, 47, boulevard de l’Hôpital, F-75013 France.

## Abstract

Lysosome dysfunction has been widely implicated in many models of neurodegeneration, but much less is understood of its involvement during brain development in health and disease. Hereditary spastic paraplegia caused by mutations in the *SPG11* gene is a neurodegenerative disorder characterized by lysosome dysfunction, which also presents neurodevelopmental alterations. Using knockout mouse and cortical organoid models derived from induced pluripotent stem cells, we show that lysosome dysfunction caused by *SPG11* mutations decreases the proliferation of neural progenitor cells at early stages of cortical development. At the cellular level, *SPG11* mutations cause accumulation of calcium in lysosomes, which reduces proliferation of neural progenitor cells and diminishes apical tight junctions. RNA sequencing analysis revealed that these phenotypes in SPG11 organoids are caused by hypoactivation of mammalian target of rapamycin (mTOR) signaling. The latter is a consequence of lysosomal recruitment of the enzyme PI4K2A (phosphatidylinositol 4-kinase type 2 alpha) resulting in higher levels of its product PI(4)P (phosphatidylinositol-4-phosphate), a described regulator of the mTOR pathway. Modulating the function of the lysosomal calcium channel TRPML1 successfully corrected all developmental phenotypes in cortical organoids, highlighting the critical role of lysosomal calcium in signaling during the early phase of cortical development.

## Introduction

Lysosomes are membrane-limited organelles that play a critical catabolic role by degrading the content of late endosomes and autophagosomes. Beyond this classical function, lysosomes have emerged as key signaling hubs that sense nutrient availability and regulate gene expression (1). Inherited mutations affecting lysosomes therefore have wider consequences that extend beyond cellular homeostasis and impair organ function. This is illustrated by lysosomal storage disorders—genetic diseases caused by deficiencies in lysosomal metabolic function that are characterized by early multisystemic alterations, frequently including central nervous system dysfunction. These disorders often impact development, with symptoms manifesting early in life (2,3). Despite this clinical evidence, the precise physiological roles of lysosomes in organ development, particularly brain development, remain poorly understood (1).

To address this question, we used hereditary spastic paraplegia type 11 (SPG11) as a disease model. SPG11 is a neurodegenerative disorder caused by mutations in the *SPG11* gene (4) that is characterized by lysosomal dysfunction (5–7) and alterations in brain development (8,9). SPG11 patients develop progressive spasticity of the lower limbs beginning in the second decade of life, often accompanied by additional neurological symptoms including ataxia, peripheral neuropathy, and cognitive impairment (10,11). Neuropathological studies of autopsy brains have revealed widespread neuronal death across multiple brain regions (12–14), confirming the neurodegenerative nature of the disease. However, mounting evidence indicates that neurodevelopmental alterations also contribute to SPG11 pathology (9). Many SPG11 patients exhibit poor academic performance that predates the onset of spasticity (15). In animal models, *Spg11* downregulation in zebrafish causes abnormal axonal branching during embryonic development, resulting in motor neuron defects (16,17). Similarly, models of neurodevelopment derived from SPG11 patient stem cells exhibit altered neural progenitor proliferation, premature neurogenesis, and disrupted signaling pathways (8,9,18). Together, these findings establish a critical role for *SPG11* during brain development.

Most mutations identified in SPG11 patients are nonsense or frameshift mutations (11) that result in loss of function of the gene product, spatacsin. Loss of spatacsin has been extensively documented to impair lysosomal function in both cellular and animal models (7,19–21). Spatacsin plays essential roles in regulating autophagic lysosome reformation — a pathway crucial for lysosome membrane recycling (6,19) — as well as lysosomal trafficking and motility (22). Consequently, spatacsin deficiency leads to autolysosome accumulation and increased intralysosomal storage of lipids, including cholesterol and gangliosides, ultimately contributing to neuronal death (5,21,23,24).

Beyond their catabolic functions, lysosomes serve as key signaling platforms, particularly for the mammalian target of rapamycin (mTOR) pathway (25), and lysosomal dynamics have recently been implicated in early cortical formation (26). However, how spatacsin loss impacts lysosome-dependent signaling pathways, and how these lysosomal dysfunctions impair brain development remain unknown.

Here, we analyzed cortical development in *Spg11* knockout mice and identified alterations during early cortical formation. We then used human cortical organoids derived from patient induced pluripotent stem cells (iPSCs) to investigate the impact of *SPG11* mutations on cortical development and to elucidate the broader role of lysosomes in neurodevelopment. Our work demonstrates that spatacsin deficiency impairs lysosomal calcium homeostasis, with significant downstream consequences for mTOR signaling and neural progenitor cell function.

## Results

### *Spg11* knockout mice exhibit defects in early cortical development associated with lysosome mislocalization

To investigate whether *Spg11* mutations affect cortical development, we first examined the organization of the motor cortex in mice at postnatal day 7 (P7), when the migration of cortical neurons is complete. We immunostained cortical sections with Cux1, Ctip2 and Tbr1, which are markers of the layers II/III, layer V and layer VI, respectively (Fig. 1A, Suppl. Fig 1A). This analysis revealed that both wild-type and *Spg11* knockout mice displayed overall normal cortical organization with well-established cortical layers. Measurement of the cortical thickness revealed no significant difference between wild-type and *Spg11* knockout mice at P7 (Fig. 1B). When investigating the main cortical layers, Cux1 and Ctip2 labeling showed no differences between *Spg11* knockout and control mice (Suppl. Fig 1A). By contrast, the number of Tbr1-positive neurons in layer VI was significantly lower in *Spg11* knockout compared to control mice (Fig. 1C).

**Figure 1.**
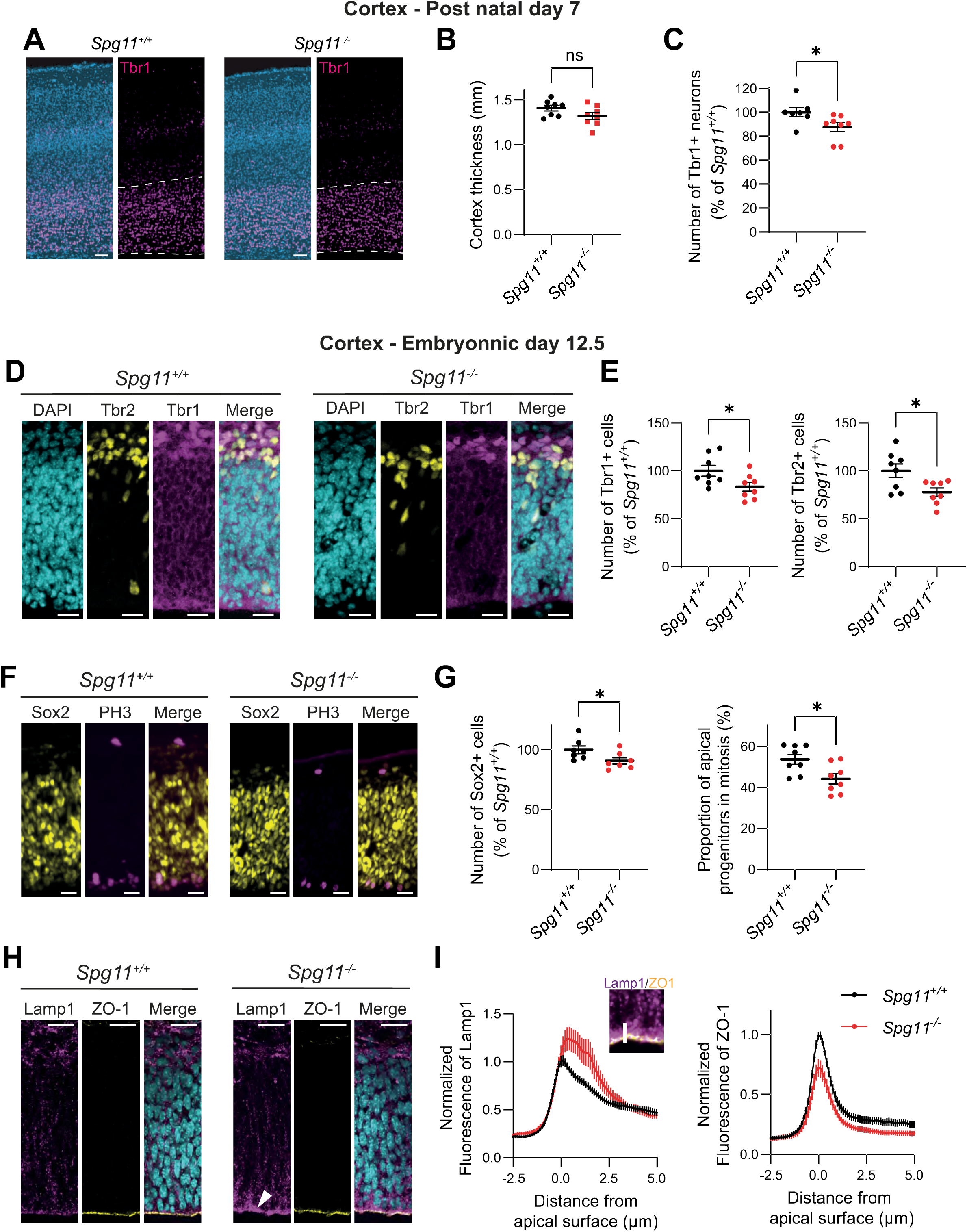
Spg11 knockout mice show impaired cortical development. A. Tbr1 immunostaining of motor cortex sections obtained from 7-day-old wild-type (*Spg11^+/+^*) or *Spg11* knockout (*Spg11^-/-^*) mice. Dashed lines indicate layer VI of cortex. Scale bar: 100 µm. B. Quantification of motor cortex thickness of wild-type (*Spg11^+/+^*) and *Spg11* knockout (*Spg11^-/-^*) mice. The graphs show the mean ± SEM. N=8 independent animals. Mann-Whitney test, ns: p>0.1. C. Quantification of the relative number of Tbr1-positive neurons in motor cortex of wild-type (*Spg11^+/+^*) or *Spg11* knockout (*Spg11^-/-^*) mice. The graphs show the mean ± SEM. N=8 independent animals. Mann-Whitney test, *p<0.05. D. Tbr1 and Tbr2 immunostaining in sections of developing cortex obtained from embryonic day 12.5 wild-type or *Spg11* knockout (*Spg11^-/-^*) mice. Scale bar: 20 µm. E. Quantification of the relative number of Tbr1-positive or Tbr2-positive cells in cortical sections obtained from embryonic day 12.5 wild-type or *Spg11* knockout (*Spg11^-/-^*) mice. The graphs show the mean ± SEM. N=8 independent animals. Mann-Whitney test, *p<0.05. F. Sox2 and phospho-histone H3 (PH3) immunostaining sections of developing cortex obtained from embryonic day 12.5 wild-type or *Spg11* knockout (*Spg11^-/-^*) mice. Scale bar: 20 µm. G. Quantification of the relative number of SOX2 positive progenitor cells (left) and the proportion of apical Sox2-positive cells that are also positive for the mitotic marker PH3 (right) in cortical sections obtained from embryonic day 12.5 wild-type or *Spg11* knockout (*Spg11^-/-^*) mice. The graphs show the mean ± SEM. N=8 independent animals. Mann-Whitney test, *p<0.05. H. Lamp1 and ZO-1 immunostaining sections of developing cortex obtained from embryonic day 12.5 wild-type or *Spg11* knockout (*Spg11^-/-^*) mice. Arrowhead indicates concentration of Lamp1 immunostaining close to the apical surface of progenitors. Scale bar: 20 µm. I. Quantification of Lamp-1 and ZO-1 fluorescence along a line scan perpendicular to the apical surface of progenitors (white line on the inset image taken from panel G; ZO-1 in yellow, Lamp1 in magenta–), indicating localization in neural progenitors of lysosome and apical junction protein. The graphs show the mean ± SEM. N=8 independent animals were analyzed.

To determine whether this deficit originates during neurogenesis, we analyzed the cortex of control and *Spg11* knockout mice at embryonic day 12.5 (E12.5) when the layer VI neurons are generated (27). Tbr1 immunostaining in the developing cortex of E12.5 mice revealed fewer Tbr1-positive neurons in *Spg11* knockout than in control embryos (Fig. 1D-E), suggesting that *Spg11* mutations impair the generation of layer VI neurons during early cortical development.

To explore the mechanisms underlying this reduction in Tbr1-positive neurons, we examined intermediate and early progenitors that contribute to the generation of these neurons, by immunostaining with the markers T-box brain protein 2 (Tbr2) and Sox2, respectively. Both the numbers of Tbr2-positive intermediate progenitors and Sox2-positive early progenitors were significantly lower in *Spg11* knockout than in control mice at this developmental stage (Fig. 1D-G), suggesting that the lower number of Tbr1-positive neurons results from impaired progenitor proliferation. To test this hypothesis, we monitored mitotic cells by immunostaining with phospho-histone H3 antibody. *Spg11* knockout embryos displayed a significantly lower proportion of mitotic progenitors compared to controls (Fig. 1F-G), confirming a lower progenitor cell proliferation.

As *Spg11* mutations impair lysosome function in various cell types (6,7,19–21), we hypothesized that lysosomal dysfunction might contribute to abnormal cortical development observed in *Spg11*-knockout animals. To test this hypothesis, we immunostained wild-type and *Spg11*-knockout E12.5 cortices with an anti-Lamp1 antibody to detect lysosomes. *Spg11*-knockout samples exhibited higher concentration of lysosomes near the apical surface of neural progenitor cells, as labelled by the tight junction protein zonula occludens-1 (ZO-1) (Fig. 1H). Quantification of the Lamp1 fluorescence intensity along a line scan perpendicular to the apical surface of the progenitors confirmed the mislocalization of lysosomes in progenitors in E12.5 *Spg11*-knockout cortices (Fig. 1I). Because the proliferation of progenitor cells depends on proper organization of their cellular junctions (28,29), we also quantified ZO-1 intensity using line scan analysis. This revealed lower ZO-1 levels at the apical surface of progenitors in *Spg11* knockout embryos compared to controls (Fig. 1H), suggesting that *Spg11* mutations disrupt the tight junction organization of neural progenitor cells.

Together, these data demonstrate that *Spg11* mutations impair early cortical development by reducing neural progenitor proliferation, which correlates with abnormal lysosomal localization and disrupted apical tight junctions in these cells.

### Lysosomal calcium homeostasis is impaired in SPG11 neural progenitors

To investigate which physiological functions of lysosomes might be impaired by *SPG11* mutations and contribute to altered neural progenitor cell proliferation, we examined human neural progenitor cells (hNPCs) derived from iPSCs. We used two isogenic hNPC lines: one derived from a healthy control subject and another in which we introduced a homozygous pathogenic nonsense mutation (c.6100C>T; p.R2034X) into the *SPG11* gene by genome editing (24).

We first assessed whether *SPG11* mutations affect the catalytic activity of lysosomal hydrolases using fluorescent reporters. Both control and SPG11 hNPCs exhibited similar lysosomal catabolic activity, as measured by the DQ-BSA unquenching assay and the Magic Red Fluorescent Cathepsin B assay (Fig. 2A, Suppl. Fig. 2A). Consistent with these findings, western blot analyses revealed comparable levels of the lysosomal protease Cathepsin D and the lysosome membrane protein LAMP1 in control and SPG11 hNPCs (Suppl. Fig. 2B-D). These results indicate that the homozygous *SPG11* mutations do not impair the catabolic activity of lysosomes in neural progenitor cells.

**Figure 2.**
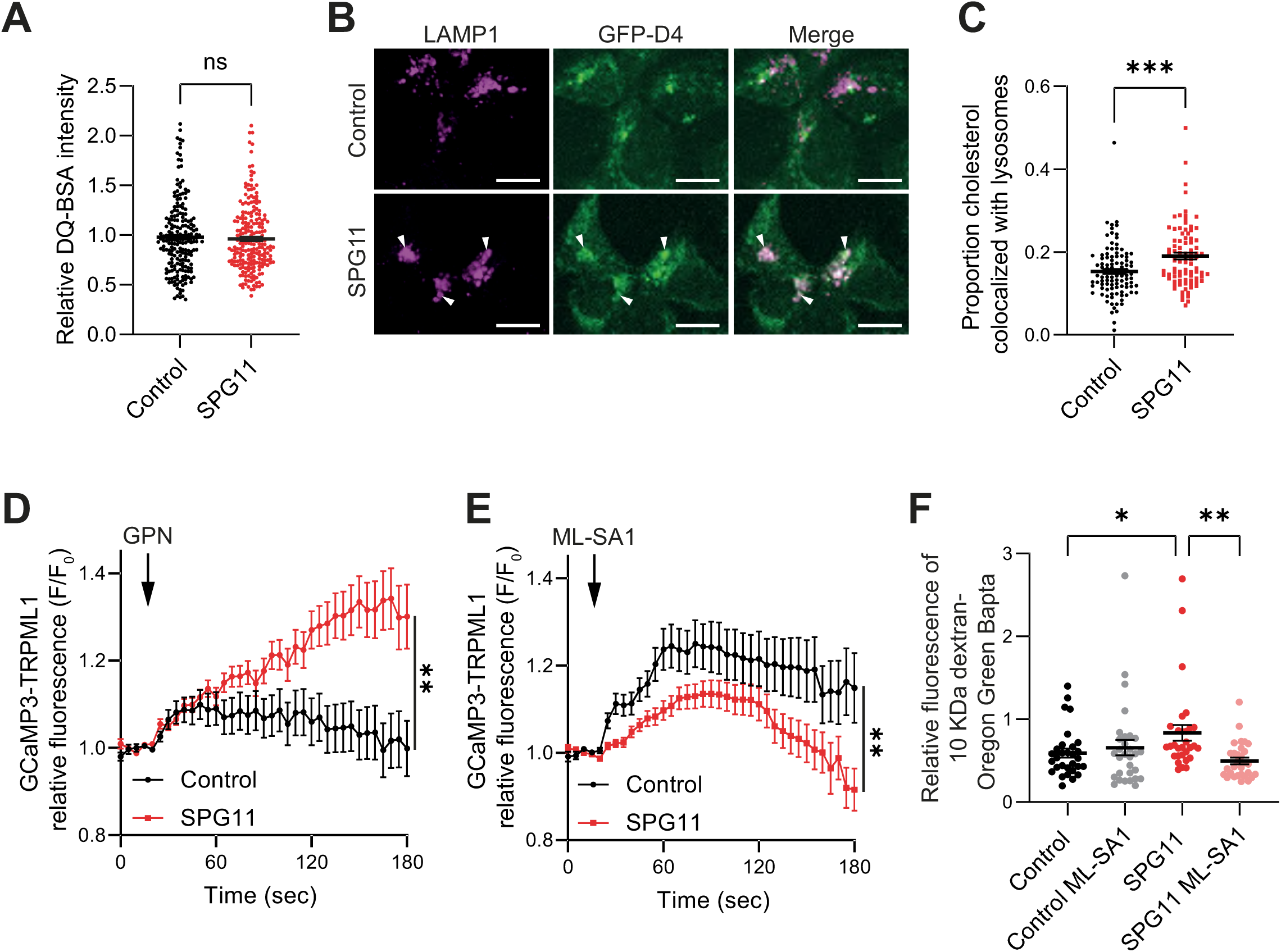
SPG11 neural progenitor cells (hNPCs) feature altered lysosomal calcium. A. Quantification of the DQ-BSA fluorescence intensity, indicative of lysosomal catalytic activity, in control and SPG11 hNPCs. The graph shows the mean ± SEM. N>200 cells from 3 independent experiments. Unpaired T-test, ns p>0.6. B. Control and SPG11 hNPCs were immunostained with a LAMP1 antibody and with the GFP-D4 probe that labels cholesterol. Arrowheads point lysosomes. Scale bar: 10µm. C. Quantification of the proportion of GFP-D4 staining colocalized with the marker LAMP1 in control and SPG11 hNPCs, showing more cholesterol in lysosomes in SPG11 hNPCs. The graph shows the mean ± SEM. N > 85 cells from three independent experiments. Unpaired t-test, *** p=0.0003. D. Relative intensity of fluorescence detected by the calcium probe GCaMP3 addressed at the surface of lysosomes (GCaMP3-TRPML1) upon osmotic shock induced by 200 µM GPN in control and SPG11 hNPCs. Osmotic shock induces the rupture of the lysosome membrane and the release of lysosomal calcium. The graph shows the mean ± SEM at each time point. N>35 cells from 3 independent experiments. Two-Way ANOVA, ** p=0.004. E. Relative intensity of fluorescence detected by the calcium probe GCaMP3 addressed at the surface of lysosomes (GCaMP3-TRPML1) upon treatment with the TRPML1 agonist ML-SA1 (20 µM) in control and SPG11 hNPCs. The graph shows the mean ± SEM at each time point. N>50 cells from 3 independent experiments. Two-Way ANOVA, ** p=0.01. F. Quantification of the relative amount of lysosomal calcium monitored using the probe Oregon Green Bapta coupled to 10 kDa dextran in control and SPG11 hNPCs, either untreated or treated for 24 hours with 20 µM ML-SA1. The graph shows the mean ± SEM. N=30 cells from 3 independent experiments. One-way ANOVA followed by Holm-Sidak’s multiple comparisons test, *p=0.021, **p=0.004.

Previous studies demonstrated that loss of spatacsin function due to *SPG11* mutations promotes accumulation of lipids, particularly cholesterol, in lysosomes of fibroblasts (23). Similarly, SPG11 hNPCs presented accumulation of cholesterol in lysosomes compared to control hNPCs (Fig. 2B,C), as detected by the cholesterol binding domain D4 of prefringolysin-O fused to GFP (GFP-D4). Since the accumulation of cholesterol in lysosomes can impair the function of the lysosomal calcium channel TRPML1 (30), we hypothesized that SPG11 hNPCs may present altered lysosomal calcium levels. To test this hypothesis, we transfected hNPCs with a vector expressing the genetically encoded calcium reporter, GCaMP3, fused with the lysosomal calcium channel TRPML1 (GCaMP3-TRPML1) (30). We quantified the relative lysosomal calcium content by triggering calcium release via osmotic rupture of the lysosomal membrane using the peptide GPN (31). SPG11 hNPCs displayed significantly higher lysosomal calcium content compared to control hNPCs (Fig. 2D). To confirm this finding with an independent method, we incubated hNPCs with the calcium-sensitive probe Oregon Green BAPTA coupled to 10 kDa dextran, along with Texas Red 10 kDa Dextran as a control of dye internalization. Following internalization by fluid phase endocytosis, both fluorescent probes localized to lysosomes (Suppl. Fig. 2E). The fluorescence intensity of Oregon Green BAPTA normalized to Texas Red fluorescence revealed a moderate but consistent increase in intralysosomal calcium in SPG11 hNPCs when compared to control cells (Fig. 2F).

To show that the change in lysosomal calcium levels in SPG11 hNPCs is due to cholesterol-induced impairment of TRPML1 channel activity (30), we triggered calcium release in GCaMP3-TRPML1 transfected hNPCs by using the TRPML1 agonist ML-SA1 (31). SPG11 hNPCs exhibited a significantly lower response to acute ML-SA1 treatment compared to control cells (Fig. 2E), suggesting that TRPML1 channel activity is reduced when *SPG11* is mutated. This dysfunction likely accounts for the higher intralysosomal calcium levels observed in these cells. Notably, a 24-hour treatment with ML-SA1 normalized lysosomal calcium levels in SPG11 hNPCs, as measured by the Oregon Green BAPTA probe (Fig. 2F).

Together, these data demonstrate that hNPCs harboring homozygous *SPG11* mutations retain normal lysosomal catabolic activity, but exhibit elevated lysosomal calcium levels due to impaired TRPML1 function. Importantly, pharmacological activation of TRPML1 with ML-SA1 can restore lysosomal calcium homeostasis in *SPG11* neural progenitors.

### Altered calcium levels contribute to lysosomal dysfunction in SPG11 cortical organoids

To investigate SPG11-induced lysosomal dysfunction in human neurodevelopment, we generated human cortical organoids and performed LAMP1 immunostaining to label lysosomal compartments. In 14-day-old SPG11 cortical organoids, lysosomes accumulated at the apical (inner) surface of neural rosettes, as labelled by ZO-1 (Fig. 3A). Quantification of the fluorescence intensity of LAMP1 and ZO-1 along a line scan perpendicular to the apical surface of the rosettes confirmed lysosomes mislocalization in SPG11 rosettes (Fig. 3B), accompanied by lower enrichment of the junction protein ZO-1 (Suppl. Fig. 3A). These findings recapitulate our observations made in the developing cortex of *Spg11* knockout mice (Fig. 1H-I).

**Figure 3.**
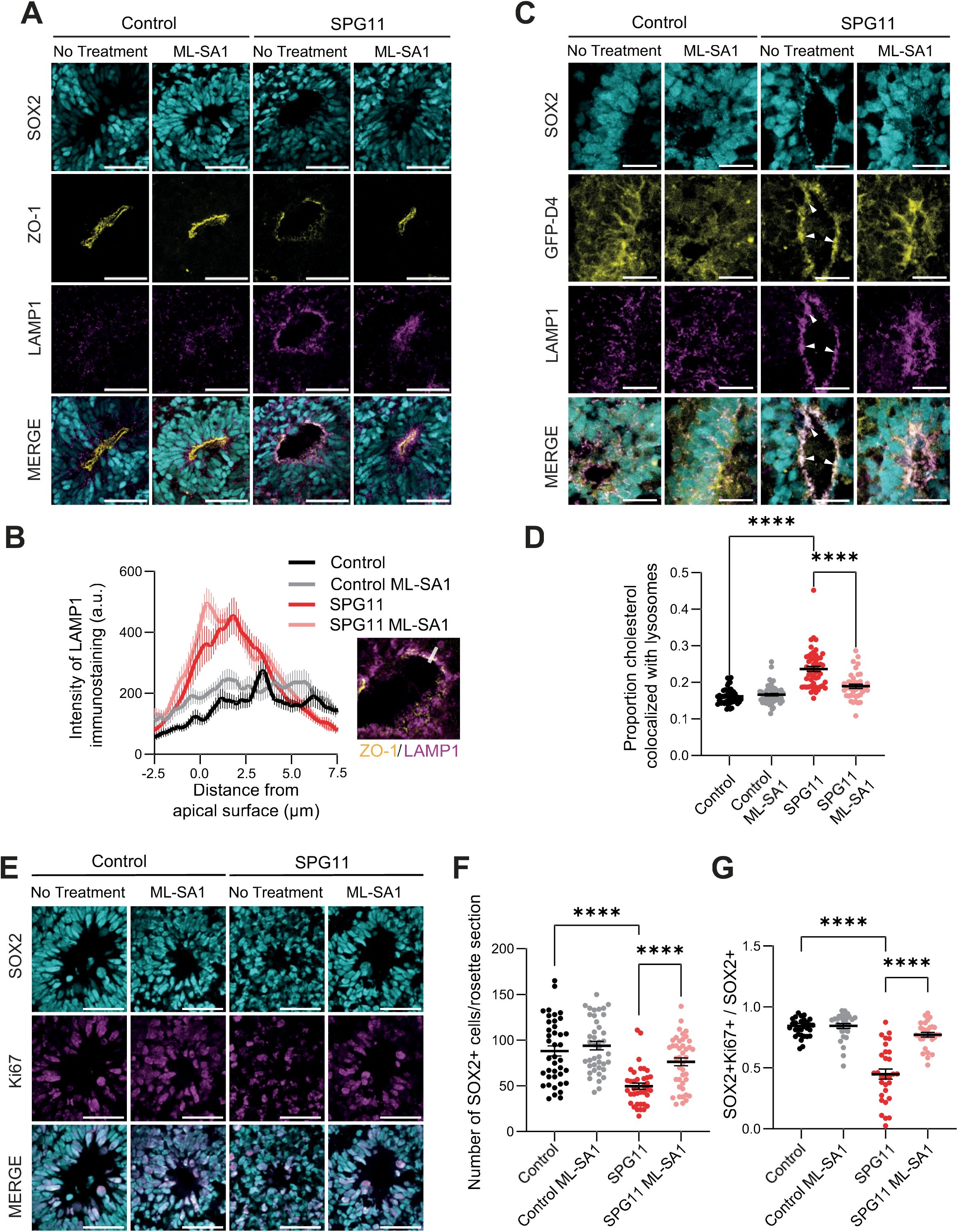
SPG11 neural progenitors in cortical organoids feature developmental impairments that are rescued by modulating lysosomal calcium channels. A. Rosettes of control and SPG11 14-day old cortical organoids, either untreated or treated with ML-SA1. Samples were immunostained with antibodies against SOX2 for neural progenitors, ZO-1 for tight junctions, and LAMP1 as a marker for lysosomes. Scale bar: 50 µm. B. Quantification of LAMP-1 fluorescence along a line scan perpendicular to the apical surface of rosettes in cortical organoids (white line on the inset image taken from panel A; ZO-1 in yellow, LAMP1 in magenta), indicating lysosome localization in neural progenitors. The graph shows the mean ± SEM. N=30 rosettes from 3 independent experiments. C. Rosettes of control and SPG11 14-day-old cortical organoids, either untreated or treated with ML-SA1, were immunostained with antibodies against SOX2 for neural progenitors as well as LAMP1 for lysosomes and were stained with the GFP-D4 probe to detect cholesterol. Note that cholesterol accumulates mainly in lysosomes in untreated SPG11 cortical organoids (arrowheads) Scale bar: 20 µm. D. Quantification of the proportion of GFP-D4 staining that is colocalized with lysosomes (LAMP1 immunostaining). The graph shows the mean ± SEM. N>40 rosettes from 3 independent experiments. One-way ANOVA followed by Holm-Sidak’s multiple comparisons test, ****p<0.0001. E. Immunostaining of rosettes in control and SPG11 14-day-old cortical organoids, either untreated or treated with ML-SA1, with antibodies against SOX2 for neural progenitors and Ki67 for dividing cells. Scale bar: 50 µm. F. Quantification of the number of SOX2 positive progenitor cells per rosette in control and SPG11 organoids that were untreated or treated with ML-SA1. The graph shows the mean ± SEM. N>35 rosettes from 3 independent experiments. One-way ANOVA followed by Holm-Sidak’s multiple comparisons test, ****p<0.0001. G. Quantification of the proportion of apical SOX2 positive cells that were also positive for the Ki67 marker in control and SPG11 organoids that were untreated or treated with ML-SA1. The graph shows the mean ± SEM. N=30 rosettes from 3 independent experiments. One-way ANOVA followed by Holm-Sidak’s multiple comparisons test, ****p<0.0001.

Given the cholesterol accumulation and impaired calcium homeostasis we observed in SPG11 hNPCs, we investigated whether these phenotypes manifested in SPG11 cortical organoids. Staining with the GFP-D4 probe revealed higher amounts of cholesterol in neural progenitors of SPG11 organoids compared to controls (Fig. 3C, Suppl. Fig. 3B). This cholesterol accumulation is predominantly localized to lysosomes in SPG11 neural progenitors (Fig. 3C,D). Notably, cholesterol also accumulated in lysosomes of cortical progenitors of *Spg11*-knockout embryos (Suppl. Fig. 3C-D), highlighting the relevance of our observation in organoids.

To determine whether lysosomal dysfunctions resulted from impaired lysosomal calcium levels, we evaluated the impact of chronic treatment with ML-SA1 in organoids. Treatment started on day 7 of culture and was continuous afterwards. While ML-SA1 did not affect the localization of lysosomes at the apical surface in control and SPG11 organoids (Fig. 3A,B), it significantly reduced lysosomal cholesterol levels in SPG11 cortical organoids (Fig. 3C,D, Suppl. Fig. 3B). These data demonstrate that altered calcium homeostasis in SPG11 organoids contributes to lysosomal cholesterol accumulation.

### Modulation of lysosomal calcium partially restores developmental defects in SPG11 neural progenitors

We next examined whether lysosomal dysfunction impairs the proliferation of neural progenitor cells in cortical organoids. Immunostaining for SOX2, a marker of neural progenitor self-renewal, revealed a marked reduction in SOX2-positive cells within SPG11 rosettes compared to controls at day 14 of differentiation (Fig. 3E-G). ML-SA1 treatment restored the number of SOX2-positive cells per rosette in SPG11 organoids to control levels (Fig. 3E,F).

Compared to control rosettes, SPG11 rosettes presented a significant decrease in the proliferation of progenitors that was assessed by counting the number of SOX2-positive cells that were also positive for the proliferation marker Ki67 (Fig. 3E,G). This result is consistent with our findings in the cortex of E12.5 *Spg11* knockout mice (Fig. 1F-G). ML-SA1 treatment rescued the proliferation defect, restoring Ki67-positive progenitors numbers to control levels in SPG11 cortical organoids (Fig. 3E,G). Moreover, ML-SA1 normalized ZO-1 levels at the apical surface of rosettes in SPG11 organoids (Fig. 3A, Suppl. Fig. 3A) suggesting restoration of cellular junctions between progenitors, which are critical for maintaining their proliferation (28,29).

To confirm that lysosomal calcium directly regulates progenitor proliferation and cellular junctions, we treated control cortical organoids with ML-SI1, an antagonist of the TRPML1 channel (34). ML-SI1 treatment significantly decreased both neural progenitor proliferation and ZO-1 levels at the apical surface of the rosettes in control organoids (Suppl. Fig. 3E-G), phenocopying the *SPG11* mutant condition.

Together, these data establish that altered lysosomal calcium homeostasis decreases neural progenitor cell proliferation in cortical organoids, and that pharmacological activation of the TRPML1 channel with ML-SA1 can compensate for this developmental dysfunction caused by *SPG11* mutations.

### Later stage SPG11 cortical organoids recover progenitor proliferation but exhibit reduced neurogenesis that is rescued by ML-SA1 treatment

We then investigated whether the consequences of *SPG11* mutations and the effects of lysosomal calcium modulation persisted during later corticogenesis. After 42 days of differentiation, SPG11 cortical organoids exhibited similar levels of ZO-1 enrichment and progenitor proliferation as control organoids, even without ML-SA1 treatment (Fig. 4A-C). These observations suggest that compensatory mechanisms arise over time in SPG11 cortical progenitors. However, despite this normalization of proliferation, the number of SOX2-positive progenitor cells per rosette remained significantly lower in SPG11 organoids compared to controls (Fig. 4D,E), likely reflecting the cumulative impact of reduced progenitor proliferation during earlier stages of cortical development. Continuous ML-SA1 treatment from day 7 onward moderately increased the number of SOX2-positive progenitors per rosette in 42-day-old SPG11 cortical organoids, although this effect did not reach statistical significance (Fig. 4D,E).

**Figure 4.**
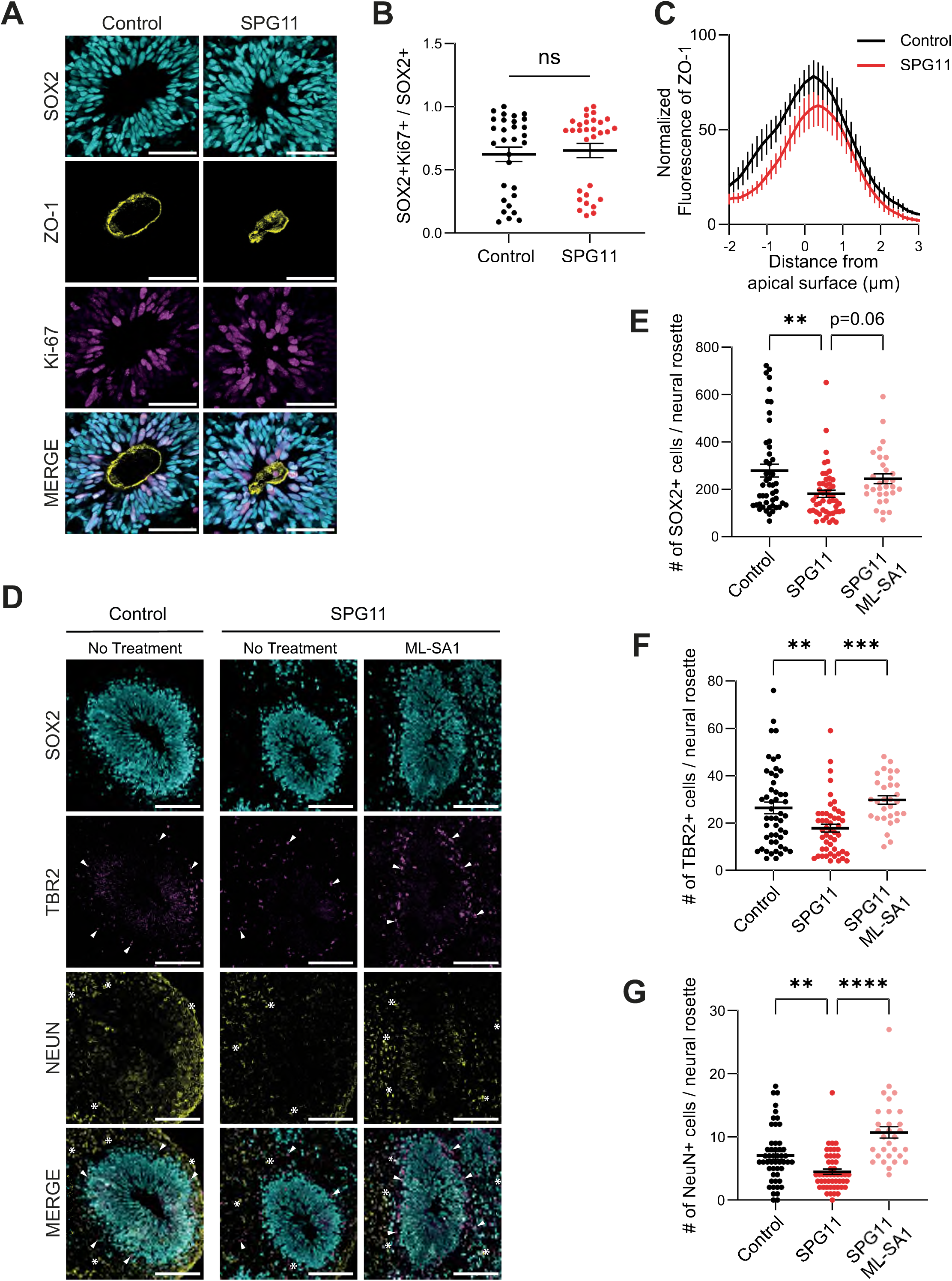
Later stage SPG11 cortical organoids show normal proliferation and tight junctions but feature reduced cortical lamination that can rescued with ML-SA1 treatment. A. Rosettes of control and SPG11 42-day-old cortical organoids were immunostained with antibodies against SOX2 for neural progenitors, ZO-1 for tight junctions, and Ki-67 as a marker for proliferation. Scale bar: 50 µm. B. Quantification of the proportion of apical SOX2 positive cells that were also positive for the Ki67 marker in 42-day-old control and SPG11 cortical organoids. The graph shows the mean ± SEM. N=30 rosettes from 3 independent experiments. Unpaired T-test, ns: p=0.69 C. Quantification of ZO-1 fluorescence along a line scan perpendicular to the apical surface of randomly selected rosettes in 42-day-old control and SPG11 cortical organoids, indicating tight junction enrichment in neural progenitors. The graph shows the mean ± SEM. N=30 rosettes from 3 independent experiments. D. Rosettes in 42-day-old control and SPG11 cortical organoids, either untreated or treated with ML-SA1, were immunostained with antibodies against SOX2 for neural progenitors, TBR2 for intermediate progenitors, and NeuN for neurons. Arrowhead point intermediate progenitors and stars point neurons. Scale bar: 100 µm. E. - G Quantification of the number of SOX2-(E), TBR1-(F) or NeuN-positive (G) cells in rosettes of 42-day-old control and SPG11 cortical organoids, either untreated or treated with ML-SA1. The graphs show the mean ± SEM. N>30 rosettes from 3 independent experiments. One-way ANOVA followed by Holm-Sidak’s multiple comparisons test, **p<0.01; ***p<0.001; ****p<0.0001.

Beyond impaired proliferation, the loss of spatacsin has been proposed to impair neurogenesis in cortical organoids (8). Control organoids at 42-days displayed characteristic laminar organization of cortical cell types, with SOX2-positive early neural progenitors being surrounded by a layer of TBR2-positive intermediate progenitors, and beyond that, NeuN-positive neurons. In contrast, SPG11 cortical organoids exhibited significantly fewer TBR2-positive intermediate progenitors and NeuN-positive neurons in laminae surrounding each rosette (Fig. 4D,F,G). ML-SA1 treatment significantly increased the numbers of both intermediate progenitors and neurons surrounding rosettes in SPG11 organoids (Fig. 4D,F,G). ML-SA1 treated control cortical organoids had poor survival at 42 days and could not be analyzed, suggesting that chronic ML-SA1 treatment may be toxic in healthy conditions.

All together, these data demonstrate that the lower number of differentiated cells surrounding rosettes in SPG11 organoids results from reduced progenitor proliferation during early corticogenesis, a defect that can be compensated by ML-SA1 treatment.

### mTOR signaling is dysfunctional in SPG11 neural progenitors and restored by ML-SA1

To investigate the molecular pathways linking lysosomal calcium homeostasis to neural progenitor proliferation, we performed bulk RNA sequencing followed by differential gene expression analysis on 14-day-old cortical organoids, when proliferation defects were most evident in SPG11 organoids. We also analyzed ML-SA1 treated cortical organoids to identify genes and pathways underlying the phenotypic improvements provided with treatment.

Comparison of SPG11 and control cortical organoids revealed 33 genes downregulated and 57 upregulated genes in SPG11 organoids (Suppl. Fig. 4A). Analysis of SPG11 cortical organoids untreated versus treated with ML-SA1 identified 14 genes downregulated and 37 upregulated genes with ML-SA1 treatment (Suppl. Fig. 4B).

To identify the pathways modified by the *SPG11* mutation or the ML-SA1 treatment, we performed gene set enrichment analysis (GSEA) that considered all genes regardless of their deregulation. Using the curated gene sets of the human Molecular Signatures Database, no pathway was identified as being significantly altered at the gene expression level between control and SPG11 organoids. In contrast, ML-SA1 treatment of SPG11 organoids only significantly modified three hallmark gene sets: mTORC1 signaling, PI3K-Akt-mTOR signaling and the p53 pathways (Suppl. Fig. 4C). Given the established link between mTOR signaling and lysosome function, and based on our findings that lysosomal dysfunction impairs progenitor cell proliferation, we focused our subsequent analysis on the impact of *SPG11* mutations and ML-SA1 on the activity of the mTOR pathway.

To assess mTOR pathway activity, we examined by immunofluorescence the phosphorylation of the ribosomal protein S6 (S6), a well-characterized downstream effector of mTOR (32). SPG11 cortical organoids featured significantly less phosphorylated S6 (pS6) within rosettes when compared to controls, and ML-SA1 treatment substantially restored pS6 levels (Fig. 5A,B). Consistent with these findings, pS6 levels were also lower in neural progenitor cells of E12.5 *Spg11* knockout embryos compared to control littermates (Suppl. Fig. 4D), highlighting the relevance of impaired mTOR signaling during early SPG11 cortical development.

**Figure 5.**
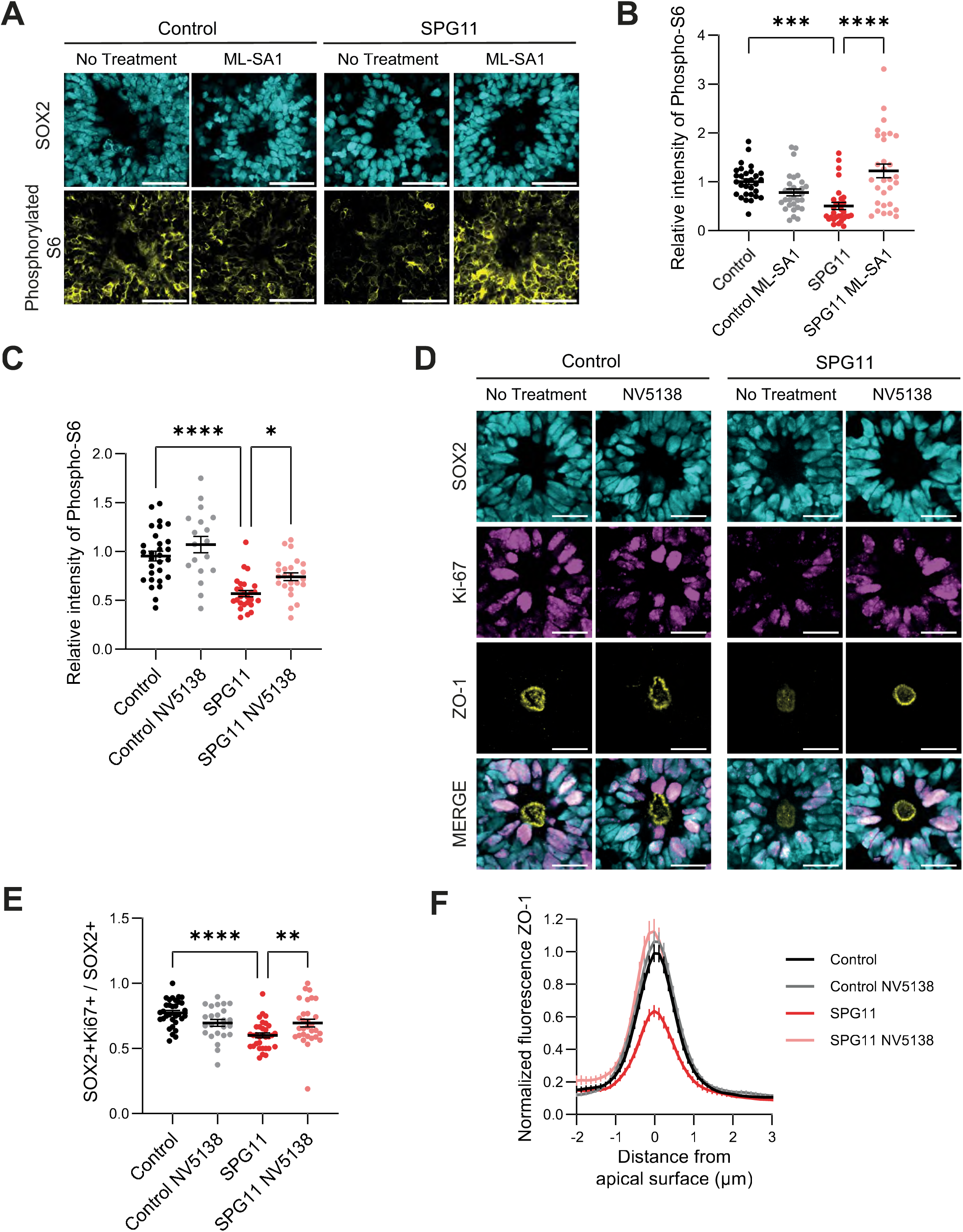
Inhibition of the mTOR pathway contributes to impaired proliferation of neural progenitors in SPG11 cortical organoids. A. Rosettes in control and SPG11 14-day-old cortical organoids, either untreated or treated with ML-SA1, were immunostained with antibodies against SOX2 for neural progenitors, and phosphorylated S6. Scale bar: 50 µm. B. Quantification of the relative levels of intensity of phosphorylated S6 in rosettes of control and SPG11 cortical organoids, either untreated or treated with ML-SA1. Values were normalized to the average fluorescence of phosphorylated S6 staining of controls in each experiment. The graph shows the mean ± SEM. N=30 rosettes from 3 independent experiments. One-way ANOVA followed by Holm-Sidak’s multiple comparisons test, ***p<0.001; ****p<0.0001. C. Quantification of the relative levels of intensity of fluorescence of phosphorylated S6 in rosettes of control and SPG11 14-day-old cortical organoids, either untreated or treated with the mTOR agonist NV5138 (100µM). Values were normalized to the average phosphorylated S6 fluorescence of controls in each experiment. The graph shows the mean ± SEM. N>25 rosettes from 3 independent experiments. One-way ANOVA followed by Holm-Sidak’s multiple comparisons test, *p=0.016;****p<0.0001. D. Immunostaining of rosettes in 14-day-old control and SPG11 organoids, either untreated or treated with NV5138, with antibodies against SOX2 for neural progenitors, KiI67 for dividing cells, and ZO-1 to label tight junctions. Scale bar: 20 µm E. Quantification of the proportion of apical SOX2 positive cells that were also positive for the Ki67 marker in control and SPG11 organoids that were untreated or treated with NV5138. The graph shows the mean ± SEM. N>25 rosettes from 3 independent experiments. One-way ANOVA followed by Holm-Sidak’s multiple comparisons test, **p=0.008; ****p<0.0001. F. Quantification of ZO-1 fluorescence along a line scan perpendicular to the apical surface of randomly selected rosettes in 14-day-old control and SPG11 cortical organoids, either untreated or treated with NV5138. The graph shows the mean ± SEM. N>35 rosettes from 3 independent experiments. Note the decrease in the intensity of ZO-1 immunostaining at the apical surface of SPG11 rosettes, which is compensated in the presence of NV5138 treatment.

To determine whether direct activation of mTOR signaling could rescue the phenotypes in SPG11 organoids, we treated organoids with the mTOR agonist NV5138 (33,34) and we monitored pS6 levels by immunofluorescence. NV5138 treatment significantly increased pS6 levels in rosettes of 14-day-old SPG11 organoids (Fig. 5C, Suppl. Fig. 4E), confirming that the treatment rescued mTOR pathway activity in SPG11 organoids. Importantly, NV5138 treatment also significantly increased neural progenitor proliferation in rosettes, as monitored by the proportion of SOX2-positive cells co-expressing the proliferation marker Ki67 in rosettes (Fig. 5D,E). Furthermore, mTOR activation with NV5138 restored normal levels of the junction protein ZO-1 in rosettes of 14-day-old SPG11 cortical organoids (Fig. 5D,F).

Altogether, these data demonstrate that the downregulation of the mTOR signaling pathway in SPG11 neural progenitors underlies both the impaired proliferation and aberrant apical junctions observed in SPG11 cortical organoids.

### ML-SA1 restores mTOR signaling in SPG11 cells by modulating lysosomal levels of phosphatidylinositol-4-phosphate

To elucidate the molecular mechanisms underlying the downregulation in mTOR signaling in SPG11 organoids, we used cultured hNPCs as a model. Consistent with our observations in organoids, SPG11 hNPCs displayed lower pS6 levels by western blot, and ML-SA1 treatment significantly increased pS6 levels in SPG11 hNPCs (Fig. 6A,B).

**Figure 6.**
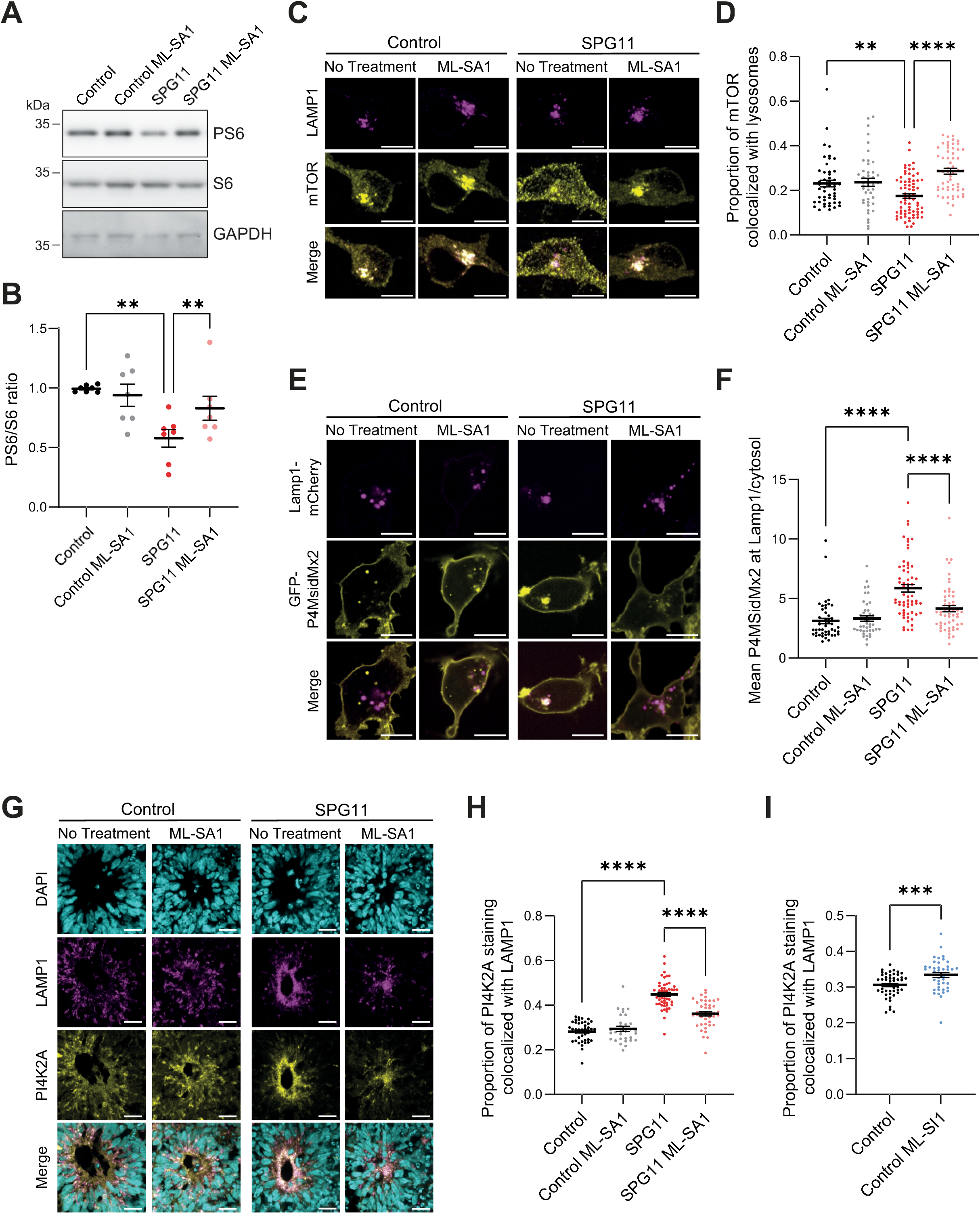
ML-SA1 restores mTOR signaling in SPG11 cells by modulating lysosomal levels of phosphatidylinositol-4-phosphate (PI(4)P). A. Western blots showing levels of phosphorylated S6 (PS6), total S6 protein, as well as GAPDH in control and SPG11 hNPCs, either untreated or treated with 20 µM ML-SA1 for 24 hours. B. Quantification of the ratio of phosphorylated to total levels of S6 (PS6/S6 ratio) in control and SPG11 hNPCs, either untreated or treated with ML-SA1. The graph shows the mean ± SEM. N=8 cell sample preparations from 5 independent experiments. One-way ANOVA followed by Holm-Sidak’s multiple comparisons test, **p<0.01. C. Immunostaining of control and SPG11 hNPCs, either untreated or treated with ML-SA1 for 24 hours, with antibodies against LAMP1 labelling lysosomes and mTOR. Scale bar: 10 µm. D. Quantification of the proportion of mTOR immunostaining that is colocalized with lysosomes (LAMP1 immunostaining). The graph shows the mean ± SEM. N> 44 transfected cells from 3 independent experiments. One-way ANOVA followed by Holm-Sidak’s multiple comparisons test, ** p<0.01; ****p<0.0001. E. Live spinning disk confocal images of control and SPG11 hNPCs, either untreated or treated with ML-SA1 for 24 hours, and transfected with vectors allowing expression of the lysosomal marker Lamp1-mCherry and the reporter of PI(4)P GFP-P4MsidMx2. Scale bar: 10 µm. F. Quantification of the GFP-P4MsidMx2 fluorescence intensity in Lamp1-mCherry mask normalized by the cytosolic levels in control and SPG11 hNPCs, either untreated or treated with ML-SA1. The graph shows the mean ± SEM. N>40 transfected cells from 3 independent experiments. One-way ANOVA followed by Holm-Sidak’s multiple comparisons test, ****p<0.0001. G. Rosettes in 14-day-old control and SPG11 cortical organoids, either untreated or treated with ML-SA1, were immunostained with antibodies against LAMP1 and PI4K2A, as well as stained with DAPI. Scale bar: 20 µm. H. Quantification of the proportion of PI4K2A colocalized with lysosomes in rosettes of control or SPG11 organoids, either untreated or treated with ML-SA1. The graph shows the mean ± SEM. N>30 rosettes from 3 independent experiments. One-way ANOVA followed by Holm-Sidak’s multiple comparisons test, ****p<0.0001. I. Quantification of the proportion of PI4K2A colocalized with lysosomes in rosettes of control organoids, either untreated or treated with ML-SI1. The graph shows the mean ± SEM. N>30 rosettes from 3 independent experiments. Unpaired T-test, ***p=0.0009.

*SPG11* mutations have previously been associated with lysosomal accumulation of phosphatidylinositol 4-kinase 2α (PI4K2A) (35). This enzyme promotes the synthesis of the signaling lipid phosphatidylinositol-4-phosphate (PI(4)P), which contributes to repress mTORC1 signaling and inhibits its localization at the lysosomal surface (36). We therefore hypothesized that ML-SA1 treatment modulates the mTOR pathway by altering phosphoinositide signaling. Colocalization experiments revealed that mTOR was less present on lysosomes in SPG11 hNPCs compared to controls, a defect that was rescued by ML-SA1 treatment (Fig. 6C,D). These findings are consistent with ML-SA1 alleviating the mTOR repression in SPG11 cells.

To determine whether altered mTOR signaling was linked to lysosomal PI(4)P levels, we expressed in hNPCs the bacterial effector protein SidM (P4MSidMx2) fused to GFP as a specific optical reporter to monitor lysosomal PI(4)P (37,38). Compared to control hNPCs, SPG11 hNPCs exhibited significantly higher PI(4)P signal at lysosomes labelled by Lamp1-mCherry (Fig. 6E,F). Importantly, ML-SA1 treatment normalized lysosomal PI(4)P levels to control values. The accumulation of PI(4)P in SPG11 hNPCs was associated with higher recruitment of the PI(4)P synthesizing enzyme PI4K2A to lysosomes, an effect that was reversed by ML-SA1 treatment (Suppl. Fig. 5A,B).

Finally, to evaluate whether this pathway contributes to the downregulation of mTOR signaling in cortical organoids and mouse embryos, we immunostained E12.5 embryos and 14-day-old organoids for PI4K2A. Consistent with observations in hNPCs, PI4K2A exhibited greater colocalization with lysosomes in neural progenitor cells of E12.5 *Spg11* knockout embryos compared to control littermates (Suppl. Fig. 5C,D), as well as in rosettes of SPG11 organoids compared to controls (Fig. 6G-H). In SPG11 organoids, this effect was normalized by ML-SA1 treatment (Fig. 6G-H). Conversely, treatment of control organoids with the TRPML1 antagonist ML-SI1 significantly increased PI4K2A localization to lysosomes (Fig. 6I), demonstrating that the lysosomal recruitment of this enzyme is regulated by lysosomal calcium.

Together, these data demonstrate that PI4K2A accumulation on lysosomes in SPG11 neural progenitor cells elevates lysosomal PI(4)P levels, thereby repressing mTOR signaling. ML-SA1 prevents PI4K2A recruitment to lysosomes by modulating lysosomal calcium release through TRPML1, thus decreasing lysosomal PI(4)P levels and restoring mTOR signaling, which rescues neural progenitor cell proliferation during early cortical development.

## Discussion

Lysosome dysfunction has been widely implicated in many models of neurodegeneration (39–41), but its role during brain development has not been properly investigated (1). Lysosomes have recently been recognized as key signaling hubs regulating cell growth, division and differentiation (42). Our study explores the role of lysosomes in regulating neural progenitor proliferation, a critical event in cortical development.

We used hereditary spastic paraplegia SPG11 as a model, since loss of function mutations in *SPG11*, encoding the protein spatacsin, cause lysosomal dysfunction (5,6,19,21) and neurodegeneration, and are associated with aberrant neurodevelopment (8,9,17,18). Analysis of *Spg11* knockout mice and cortical organoids derived from *SPG11*-mutated iPSCs revealed that spatacsin loss of function impairs the early stages of cortical development and affects the formation of the early-generated neurons. In both models, *SPG11* mutations reduced neural progenitor proliferation during early cortical development, which was associated with accumulation of lysosomes close to the apical surface of progenitors and reduced activation of the mTOR pathway. The lower proliferation of progenitors in early-stage organoids resembles neurodevelopmental phenotypes observed in microcephaly models (43,44). However, this phenotype is transient, as later-stage organoids show normal proliferation of apical progenitors, and *Spg11* knockout mice exhibit lower number of neurons only in early-born cortical layer VI, whereas other layers are preserved. These findings suggest that compensatory mechanisms operate during later stages of cortical development.

Spatacsin loss of function has been associated with several lysosomal dysfunctions, including lipid accumulation in lysosomes (5,21,23) and impaired lysosomal trafficking (22). Loss of spatacsin also disrupts autophagic lysosome reformation, leading to autolysosome accumulation (6,19), although the catalytic activity of lysosomal enzymes appears to be preserved (6). Here we identify a slight yet consistent increase in lysosomal calcium levels in cultured SPG11 hNPCs, which can be rescued by a treatment with ML-SA1, an agonist of the lysosomal calcium channel TRPML1. The normalization of cholesterol or phosphoinositide levels in lysosomes of SPG11 cortical organoids following ML-SA1 treatment suggests that impaired lysosomal calcium homeostasis contributes to dysfunction in SPG11 cortical organoids. Studies in cellular models of Niemann Pick disease type C, which present accumulation of cholesterol, have suggested that a reduced TRPML1-mediated lysosomal calcium release (30) leads to higher lysosomal calcium levels. We also observed dysregulation of lysosomal phosphoinositide levels in SPG11 hNPCs. Since the lysosomal calcium channels, such as TRPML1 or TPC2, are regulated by PI(3,5)P2 levels (45), phosphoinositide dysregulation may sustain or contribute to impaired lysosomal calcium homeostasis. Alternatively, other dysregulated mechanisms could impact lysosomal calcium homeostasis (46), such as endoplasmic reticulum-lysosome contacts, which are altered in SPG11 (22) and are likely impaired in SPG11 neural progenitor cells due to abnormal lysosome localization.

In cortical organoids, an established model of early cortical development (47), we demonstrate that both neural progenitor proliferation and cellular junctions are impaired in SPG11 organoids after 14 days of differentiation. Both phenotypes can be rescued by activating the TRPML1 channel. TRPML1-mediated calcium release is impaired in several lysosomal storage disorders such as Niemann Pick diseases types A and C, Fabry disease, mucolipidosis type IV and Neuronal Ceroid Lipofuscinosis type 3 (48,49), which are each characterized by early onset and possible alterations in brain development (50–52). Neural progenitor proliferation is impaired in some models of these lysosomal storage diseases (53), but whether this dysfunction relates to impaired lysosomal calcium homeostasis remains to be investigated. Calcium levels, particularly intracellular calcium dynamics, play a central role in regulating neural progenitor cell proliferation (54). Our work highlights the critical role of lysosomal calcium in this process.

Altered lysosomal calcium homeostasis has been implicated in human diseases, where it likely influences cellular signaling (46). Our transcriptomic analysis, corroborated by immunohistochemistry analysis in cortical organoids and mouse embryonic cortices, revealed that mTOR signaling is inhibited in SPG11 models and that the TRPML1 agonist corrects the SPG11 phenotypes by activating the mTOR pathway. The latter has long been associated with impaired cortical development. Indeed, a group of neurodevelopmental disorders known as mTORopathies, often presenting as cortical dysplasia associated with epilepsy, are caused by increased mTOR signaling activity (55–57). However, hypoactivation of mTOR has also been reported to cause abnormal cortical development (58). Notably, pharmacological or genetic inhibition of mTOR impairs neural progenitor proliferation (26,59,60), similar to our observations in SPG11 models. The involvement of mTOR in SPG11 is further confirmed by the rescue of neural progenitor proliferation upon treatment of SPG11 cortical organoids with the mTOR agonist NV5138 (33,34,58).

mTOR activity is regulated by numerous growth factors and signaling pathways. In SPG11 cortical organoids, mTOR hypoactivity is corrected by treatment with the TRPML1 agonist ML-SA1, suggesting that this decrease results from impaired calcium release from lysosomes. Previous studies have similarly proposed that TRPML1-mediated lysosomal calcium release is necessary for mTORC1 activity (61,62). Loss of function mutations in *SPG11* also cause lysosomal accumulation of the enzyme PI4K2A (35). This enzyme promotes synthesis of the signaling lipid PI(4)P, which contributes to mTORC1 signal repression (36). Our study demonstrates that TRPML1 activation decreases the amount of PI4K2A localized at lysosomes in cultured neural progenitors and in cortical organoids. Since PI4K2A is inhibited by calcium (63), activation of the TRPML1 channel likely reduces PI(4)P synthesis at lysosomal membranes, thereby alleviating mTOR signaling repression. Calcium regulation of mTOR signaling appears to be a key regulatory mechanism (64), and our work highlights the importance of lysosomal calcium in this process.

In conclusion, our study establishes a clear link between lysosomal dysfunction and early cortical development and highlights its relevance using *Spg11*-knockout mouse model and SPG11 cortical organoids. Our data demonstrate that the activity of the TRPML1 lysosomal calcium channel plays a critical role in regulating neural progenitor proliferation during early phases of cortical development, by regulating the PI4K2A/mTOR pathway. These findings reveal a previously unidentified link between lysosomal calcium dynamics and early cortical development, with implications on our understanding of both neurodevelopment and neurodegeneration. Knowing that progenitor proliferation defects can be rescued by modulating TRPML1 channel activity, this work opens the possibility of evaluating in mouse models whether alterations occurring during brain development contribute to the adult phenotype observed in SPG11 patients.

## Methods

### Ethical Approval

The care and treatment of animals were in accordance with European Union Directive 2010/63/EU and with national authority (Ministère de l’Agriculture, France) guidelines for the detention, use, and ethical treatment of laboratory animals. All the experiments were approved by the local ethics committee (APAFIS reference #201604201549915), and experiments were conducted by authorized personnel in specific pathogen–free animal facilities.

The use of patient-derived material was obtained following procedures sanctioned by the ethics committee, with written and informed consent acquired from patients, as per approval of SPATAX n° DC-2008-236 - RBM029 dated 09/02/2004.

### iPSCs culture

Isogenic iPSCs were generated by introducing a homozygous stop mutation in *SPG11* (c.6100C > T, p.R2034X) in an iPSC line derived from an healthy subject (24). iPSCs were cultured in mTeSR+ (STEMCELL #100-0276) medium, on Geltrex basement membrane (Gibco A1413302), at 37°C and 5% CO_2_ with medium changes every 2-3 days. Cells were passaged using ReLeSR (STEMCELL #100-0483) every 4-7 days. iPSCs were stored for extended periods in liquid nitrogen using CryoStor (Sigma-Aldrich C3124) according to the manufacturer’s guidelines.

### Cortical Organoid Generation

Cortical organoids were generated from human iPSCs using a protocol adapted from feeder-free, xeno-free generation of cortical spheroids from human pluripotent stem cells (47). iPSCs were detached using Gentle Cell Dissociation Reagent (STEMCELL #100-0485) for 3 minutes at 37°C, and were resuspended in 1 mL of mTeSR+ medium supplemented with 10 μM Y-27632 (+Y27) (STEMCELL #72304). The appropriate volume of cell suspension was added to the AggreWell plate to achieve a density of 3.10^6^ cells per well. The plate was centrifuged at 100 x g for 1 minute, then reversed and centrifuged again at 100 x g for another 2 minutes. The plate was then incubated for 24 hours at 37°C in a 5% CO_2_ incubator. On the next day, medium was removed from the AggreWell plate without disturbing the spheroids. The iPSC-derived spheroids were harvested from the microwells by pipetting TeSR-E6 (E6) (STEMCELL #05946) medium for neural induction supplemented with SMAD inhibitors (2.5 μM Dorsomorphin [DM][STEMCELL #72102] and 10 μM SB-431542 [SB][STEMCELL #72232]) into the well. Spheroids were collected and transferred to a non-treated culture plate. The dish of cortical spheroids was then completed to a total volume of 10 mL of E6+DM+SB.

From then onwards, daily media changes were performed by removing 80-90% of the medium and replacing it with fresh medium. After 6 days, E6 medium containing DM and SB was replaced with neural medium (NM) consisting of Neurobasal-A Medium [Gibco #10888022] supplemented with B-27 minus vitamin A [1:50][Gibco #12587010], GlutaMAX [1:100][Gibco #35050038], and Antibiotic-Antimycotic [1:1000][Gibco #15240062], and further supplemented with EGF (20 ng/ml)(STEMCELL #78006) and FGF2 (20 ng/ml)(STEMCELL #78003). This medium was changed daily for the first 7 days and then every other day for the subsequent 14 days. Once organoids were 28 days old, to promote differentiation of the progenitors, FGF2 and EGF were replaced with 20 ng/mL BDNF (STEMCELL #78005) and 20 ng/mL NT-3 (STEMCELL #78074), with medium changes every other day until fixation. Treatment with ML-SA1 (TOCRIS 4746) was started at day 7 at a concentration of 20 μM and was continuous until fixation. Treatment with ML-SI1 (Sigma-Aldrich G1421) was started at day 7 at a concentration of 2 μM and was continuous until organoids were used for analysis. Treatment with NV5138 (Medchem Express HY-114384) was continuous at a concentration of 100µM from day 9 until fixation of organoids at day 14.

### Generation of hNPCs

hNPC culture was prepared by plating a dozen 14-day-old cortical organoids into a well of a Geltrex-coated 6-well plate. The medium in the well was replenished to 2 mL using N2B27 medium (50% Neurobasal-A Medium with 50% DMEM/F-12, GlutaMAX, supplemented with 1x N-2 Supplement (Gibco 17502048), 1x B-27 supplement, w/o vitamin A, and 50 μM 2-mercaptoethanol (Gibco 31350010)), completed with EGF2 (10 ng/ml), FGF2 (10 ng/ml), and BDNF (20 ng/ml). Daily medium changes with complete N2B27 medium were performed for 3 days until distinct neural rosettes emerged. To isolate neural rosettes, cells were rinsed with warm DMEM-F12, followed by the addition of 1 mL of STEMdiff Neural Rosette Selection Reagent (STEMCELL #05832). After a 90-minute incubation at 37°C, the selection reagent was removed and DMEM-F12 was pipetted firmly onto the neural rosettes to dislodge them. The medium containing the dislodged rosette clusters was collected in a tube, centrifuged at 200 x g for 5 minutes. The pellet was resuspended in complete N2B27 medium, ensuring minimal disruption to the rosettes, before being plated onto one well of a Geltrex-coated 6-well plate.

Daily medium changes were continued using complete N2B27 medium for 3-5 days until the rosette culture reached confluence. The passage process involved rinsing cells once with warm DMEM-F12, treating them with ACCUTASE (STEMCELL #07920) for 5 minutes, and subsequently detaching and neutralizing the detachment solution with warm N2B27 medium. After centrifugation at 180 x g for 4 minutes, the cells were resuspended in N2B27 complete medium and seeded onto Geltrex-coated cultureware at a density of 150,000 cells/cm^2^. hNPC identity was confirmed via immunofluorescence of markers PAX6, SOX-2, and Nestin. hNPCs were used until they reached passage 10.

### Immunostaining

Brains of E12.5 mice were dissected in 0.1 M PBS (pH 7.4) and fixed with 4% paraformaldehyde (PFA) solution in PBS at 4°C overnight. Brains of *Spg11^+/+^* and *Spg11^-/-^*embryos from the same litter were paired for analyses. For postnatal brains, 7-day-old mice were killed by decapitation and brains were dissected in PBS and fixed in 4%PFA diluted in PBS. Fixed brains were rinsed 3 times 10 minutes with PBS and were cryoprotected overnight in 30% sucrose (Sigma-Aldrich S0389) in PBS at 4°C, embedded in OCT Compound (VWR International) before being stored at -80°C. Brains were cryosectioned (14µm for E12.5 brains, 16µm for P7 brains) on a cryostat (LEICA_CM3050S) and mounted onto slides for analyses (SuperFrost-plus, Thermo-scientific) that were stored at -80°C.

At specified time points, organoids were fixed in 4% PFA diluted in PBS for 24 hours, then cryoconserved by submerging 24 hours in 30% sucrose, and embedded in Neg-50 Frozen Section Medium (Epredia 6502) before being stored at -80°C. Organoid slices (12 μm) were cut on a cryostat and mounted on Superfrost slides (Epredia J1800AMNZ) before being stored at - 80°C.

To perform immunostaining, slides were thawed for 30 minutes, and rehydrated in 1x PBS (EUROMEDEX ET330A) for 5 minutes. Blocking was performed with 1x PBS, Triton X-100 0.1% (Sigma-Aldrich T9284), and 3% BSA (EUROMEDEX ET330A 04-100-812) for 1 hour. Primary antibody solution was prepared in the same solution as during blocking and applied to slides overnight at 4°C. After three washes with PBS 1x, secondary antibodies, also in blocking solution, were applied for 1 hour. Slides were washed three times with PBS, and mounted on glass slides with Fluoromount (Sigma-Aldrich F4680).

To perform immunostaining on hNPCs, cells were fixed with PFA 4% diluted in PBS for 30 minutes. Immunostaining of hNPCs was performed as with organoids, except initial blocking was 30 minutes, and primary antibody incubation was 1 hour.

### Transcriptomic analysis of organoids

RNA was purified from fresh 14-day-old cortical organoids using Maxwell RSC simplyRNA Cells Kit (Promega AS1390) for the Maxwell RSC Instrument. mRNA library preparation was realized following manufacturer’s recommendations (ILLUMINA mRNA prep). Final samples pooled library preparations were sequenced on Nextseq 2000 ILLUMINA with P2-200 cycles cartridge (2x400Millions of 100 bases reads), corresponding to 2 x 25 million reads per sample after demultiplexing. Raw data was processed with Dragen Illumina DRAGEN bio-IT Platform (v3.10.11), using default parameters, for adapter trimming, alignment to the hg38 reference genome with the gencode v37 gtf file and expression quantification. Library orientation, library composition and coverage along transcripts were checked with Picard tools.

Downstream analyses were conducted with R (v4.1.1) via an interface provided by the Data Analysis Core of the Paris Brain Institute. Cpm-normalization was performed with edgeR (v3.28.0) for initial filtering (cpm > 1 in at least 1 sample). Differential analysis was performed with EdgeR, with thresholds on log2 fold change > 1 and FDR < 0.05. Experiments launched separately were added as a covariate to account for experimental variation. P-values were adjusted with the Benjamini-Hochberg procedure to control for multiple hypotheses testing. Enrichment analysis was conducted with clusterProfiler R package (v4.2.2) (65) with Gene Set Enrichment Analysis, on *a priori* identified pathways from the HALLMARK database.

### Antibodies and Fluorescent Molecules

Antibodies used include: anti-SOX2 (Invitrogen 14-9811-82), anti-ZO-1 (Cell Signaling Technology #13663), mouse anti-LAMP-1 (Santa Cruz sc-18821, for human samples), rabbit anti-LAMP1 (Cell Signaling Technology #9091), rat anti-Lamp1(Developmental Studies Hybridoma Bank, clone 1D4B, for mouse samples), anti-Ki-67 (BD 550609), anti-NeuN (Sigma-Aldrich MAB377), anti-Tbr1 (Abcam #ab31940), anti Ctip2 (Abcam #ab18465), anti-Cux1 (Santa Cruz Biotechnologies #sc-13024), anti-Tbr2 (Sigma-Aldrich AB2283), anti-Phospho-Histone H3 (Cell Signaling Technology #9706), anti-Phospho-S6 Ribosomal Protein (Ser240/244) (Cell Signaling Technology #5364), anti-S6 Ribosomal Protein (Cell Signaling Technology #2317), anti-PI4K2A (Santa Cruz Biotechnology sc-390026), anti-mTOR (Cell Signaling Technology #2983), anti-Cathepsin D (Abcam #ab75852), anti-β-tubulin (Abcam #ab6046) and anti-GAPDH (Sigma-Aldrich #G9545). To stain nuclei we used DAPI dilactate (Sigma-Aldrich D9564).

### Plasmids

The vector expressing the calcium probe GCaMP3 fused to TRPML1 (GCaMP3-TRPML1) (33) was obtained from Haoxing Xu (University of Michigan, USA). GFP-P4M-SidMx2 (#51472), Lamp1-mCherry (#45147) were obtained from Addgene. Plasmid allowing production of the GFP-D4 probe (66) was a gift from T. Kobayashi (CNRS, Illkrich, France).

### Western Blot

Cells were detached with ACCUTASE, washed once with PBS, and suspended in lysis buffer (100mM NaCl, 10 mM TRIS pH 7.4, 2 mM MgCl_2_, 1% SDS, 0.1% Benzonase). The suspension was then centrifuged at 17,000g for 15 minutes at room temperature, and the supernatant was collected. 20 µg of protein were loaded and separated on NuPAGE 4%-12% gels (Invitrogen NP0321) in NuPAGE MOPS (Invitrogen NP0001) buffer. Protein was transferred to Immobilon-PVDF (Millipore IPVH00010) membranes. Membranes were saturated with 5% semi-skimmed milk diluted in PBS-0.5% Tween20 for 30 minutes, followed by incubation with primary antibodies in the same solution overnight at 4°C. The following day, the membrane was washed and incubated with secondary antibodies coupled to HRP (The Jackson Laboratory) in milk for 1 hour before being imaged using SuperSignal West Femto (Thermofisher 34075/34094) with a ChemiDoc Touch system (Bio-Rad).

### Lysosomal Calcium Assay

To monitor lysosomal calcium, hNPCs were grown in Nunc Lab-TekII chambers (Thermo Scientific 171080) and were transfected with a vector expressing the calcium probe GCaMP3 fused to TRPML1 that preferentially detects juxta-lysosomal Ca^2+^ increases (30). 10^6^ cells were transfected with 5 µg of plasmid using the Neon electroporation system (Invitrogen NEON) with 3 pulses of 10 ms at 1300V according to manufacturer’s instructions, and were analyzed 24 hours later. To induce the release of calcium from lysosomes, we performed live imaging in a thermostated chamber with a Leica DMi8 inverted microscope equipped with 63× objective N.A. 1.40, a Yokogawa Confocal Spinning Disk module and a Hamamatsu Orca flash 4.0 camera. We measured changes in the fluorescence of the GCaMP3-TRPML1 probe upon addition of 200 µM GPN (Abcam, ab145914) which induces the release of lysosomal calcium upon osmotic shock causing lysosome membrane permeability, or upon addition of 20 µM ML-SA1 which induces the release of calcium through the TRPML1 channel.

As an alternative method to monitor lysosomal calcium, hNPCs grown on 8-chamber Ibidi coverglass at a density of 50,000 cells/cm2 were incubated with 0.1 mg/mL of 10 kDa dextran coupled to calcium-sensitive Oregon Green 488 BAPTA-1 (Invitrogen O6798) and 0.1 mg/mL 10 kDa Dextran coupled to Texas Red (Invitrogen D1830) overnight to allow the dyes to reach late endosomes and lysosomes upon endocytosis. The following day, the medium was changed 6 hours before analysis to clear out the excess dye. Thirty minutes before analysis, the medium was replaced with DMEM without phenol red (Gibco A1443001) supplemented either with 50 nM CellTracker Blue (Invitrogen C2110) for 30 minutes to stain the cytoplasm, or with 50nM Lysotracker blue DND-22 (Invitrogen L7525) for 30 minutes to stain lysosomes. After a wash with DMEM without phenol red, we performed live imaging at 37°C and 5% CO_2_ with a Leica DMi8 inverted microscope equipped with 63× objective N.A. 1.40, a Yokogawa Confocal Spinning Disk module and a Hamamatsu Orca flash 4.0 camera. Lysosomal calcium was assessed by normalizing the fluorescence value of calcium-sensitive BAPTA-1 Oregon Green to the fluorescence value of Texas Red in randomly selected cells.

### Lysosomal catalytic activity assays

Lysosomal catalytic activity was assessed in control and SPG11 hNPCs using two independent assays, the magic red fluorescent cathepsin B and the DQ-BSA assay. hNPCs were incubated with CellTracker™ Blue at 37°C for 30 minutes. Next, cells were incubated in medium containing Magic Red Fluorescent Cathepsin B (ImmunoChemistry Technologies 937) for 60 minutes. Finally, cells were rinsed with PBS once, fixed with 4% PFA, and then imaged with an automated ArrayScan XTI (ThermoScientific) using the General Intensity Measurement protocol, analyzing > 500 cells per well and per set of conditions.

For the second assay, cells were incubated with 5µg/ml DQ Green BSA (Thermofisher) for 1 hour at 37°C. Cells were then fixed and imaged using a with a Leica DMi8 inverted microscope equipped with 63× objective N.A. 1.40, a Yokogawa Confocal Spinning Disk module and a Hamamatsu Orca flash 4.0 camera. Signal intensity was monitored using FiJi.

### Cholesterol staining

Cholesterol was stained using a preparation of domain D4 of prefringolysin O fused to GFP (GFP-D4) that was produced and purified as previously described (66). Cells or organoid slices were first processed for immunostaining with LAMP1 antibody. Labeling of cholesterol was performed by incubating samples with 20 µg/ml recombinant GFP-D4 for 20 min at 22 °C.

### Image Analysis

Image analysis was performed using FiJi. Proliferation of progenitors was quantified by counting the number of Ki-67 positive and Sox2 positive nuclei within the first layer of cells around the rosette apical surface (as indicated by ZO-1), and dividing by the total number of SOX2 positive nuclei in the same layer of cells around the rosette apical surface. Neurogenesis was assessed by counting the number of Tbr2 and NeuN positive nuclei within the vicinity of each rosette.

To monitor enrichment of tight junction marker and lysosomes at the apical surface of organoid rosettes, we measured the fluorescence intensity of the immunostaining of ZO-1 and LAMP1, respectively, across a 3 μm-wide line perpendicular to the apical surface using the Plot profile tool in FiJi. Values were normalized to the average minimum and maximum values in control organoids of each experiment.

Fluorescence intensity of cells stained with DQ-BSA, Oregon Green 488 BAPTA-1-dextran, Texas Red-dextran and GFP-D4 (cholesterol levels), or transfected with the GCaMP3-TRPML1 probe were quantified as the mean gray value using FiJi. Colocalization of cholesterol staining (GFP-D4) with lysosomes was quantified using FiJi on images of randomly selected cells or rosettes as previously described (23). First, we created a mask corresponding to LAMP1 staining in each rosette using the IJ_IsoData threshold in FiJi. The mask was copied to the corresponding fluorescence image of cholesterol. We quantified the total intensity of cholesterol fluorescence in the lysosome mask and expressed it as the percentage of total cholesterol fluorescence in every cell or rosette. The same method was applied to mTOR and PI4K2A immunostainings.

Intensity of bands in western blot was quantified using the Gel tool in FiJi.

### Statistics

With the exception of the previously described transcriptomic analyses, all statistics were performed using GraphPad Prism version 10.2.0 (https://www.graphpad.com/). Tests performed are described in each figure description.

## Data availability

The RNA-seq datasets supporting results of this paper have been deposited at GEO, under the accession number GSE305269.

## Acknowledgments

This work benefited from equipment and services of the iGenSeq (genotyping and sequencing), ICV-iPS (cell culture), HISTOMICS (histology), ICM.Quant (microscopy; RRID: SCR_026393) and Data Analysis Core (DAC) facilities at the Paris Brain Institute (ICM). We gratefully acknowledge Alexandra Durr for providing cells used for iPSCs reprogrammation, Marie Coutelier and Simang Champramary for their assistance with RNAseq data analysis, and Sandrine Humbert for her critical comments on the manuscript. This work was supported by the “Investissements d’avenir” program grants (ANR-10-IAIHU-06) and (ANR-11-INBS-0011). The project was also supported by a FRM Equipe funding (EQU202203014671). D.S. was supported by a grant from the ED3C doctoral School (Sorbonne Université).

## Declaration of generative AI and AI-assisted technologies in the writing process

During the creation of this work, the authors utilized Claude-Sonnet 4.5 to rephrase english sentences and verify english grammar. After using this tool, the authors reviewed and edited the content as needed and take full responsibility for the content of the publication.

## Declaration of interests

Authors have no competing interests to declare.

---

**Supplementary Figure 1.**
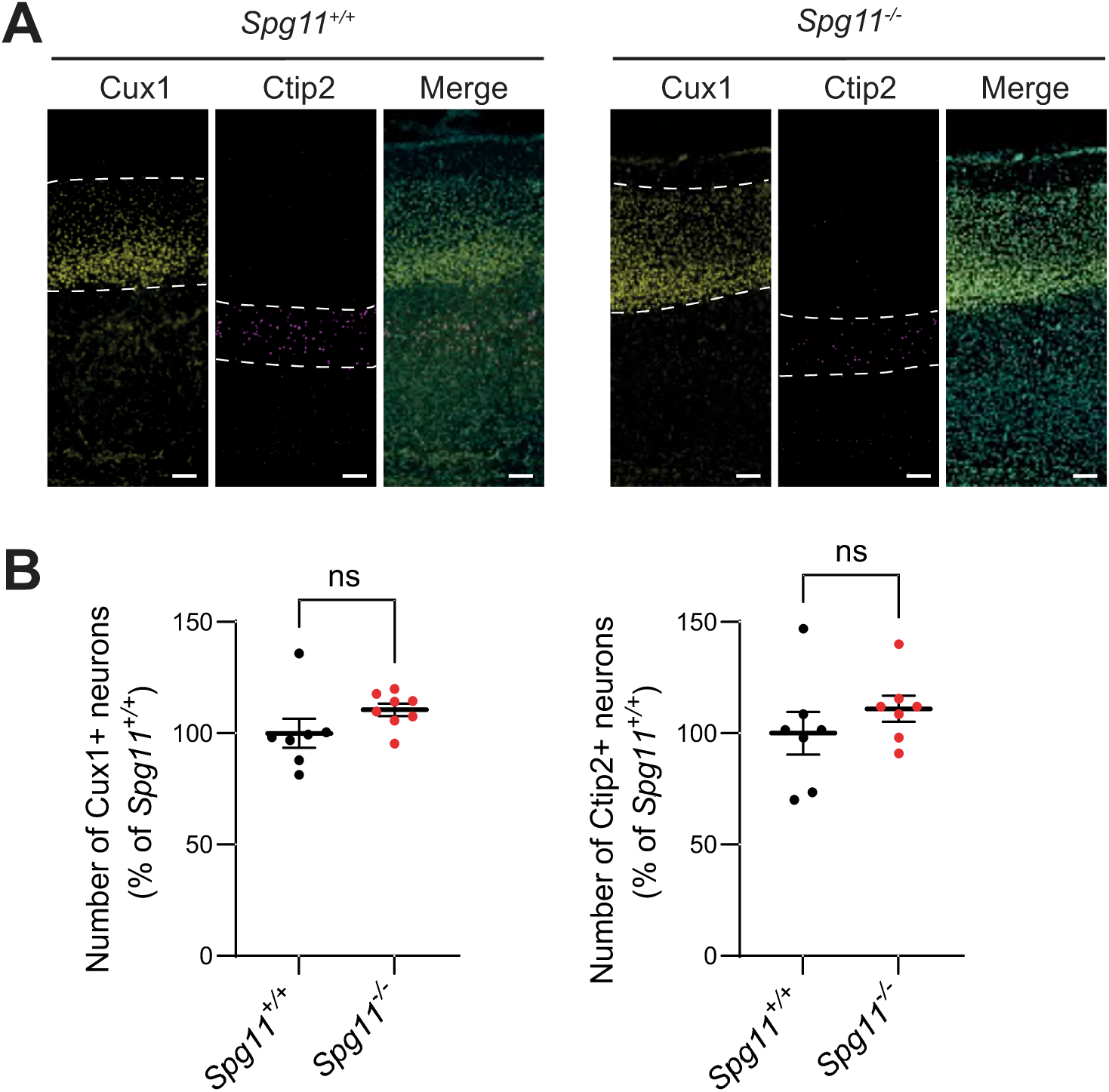
Upper cortical layers are not affected in 7-day-old *Spg11* knockout mice. A. Cux1 and Ctip2 immunostaining of cortical sections obtained from 7-day-old wild-type (*Spg11^+/+^*) or *Spg11* knockout (*Spg11^-/-^*) mice. Scale bar: 100 µm. B. Quantification of the relative number of Cux1-positive or Ctip2-positive cells in cortical sections obtained from embryonic day 12.5 wild-type (*Spg11^+/+^*) or *Spg11* knockout (*Spg11^-/-^*) mice. The graphs show the mean ± SEM. N=8 independent animals. Mann-Whitney test. ns: p>0.1.

**Supplementary Figure 2.**
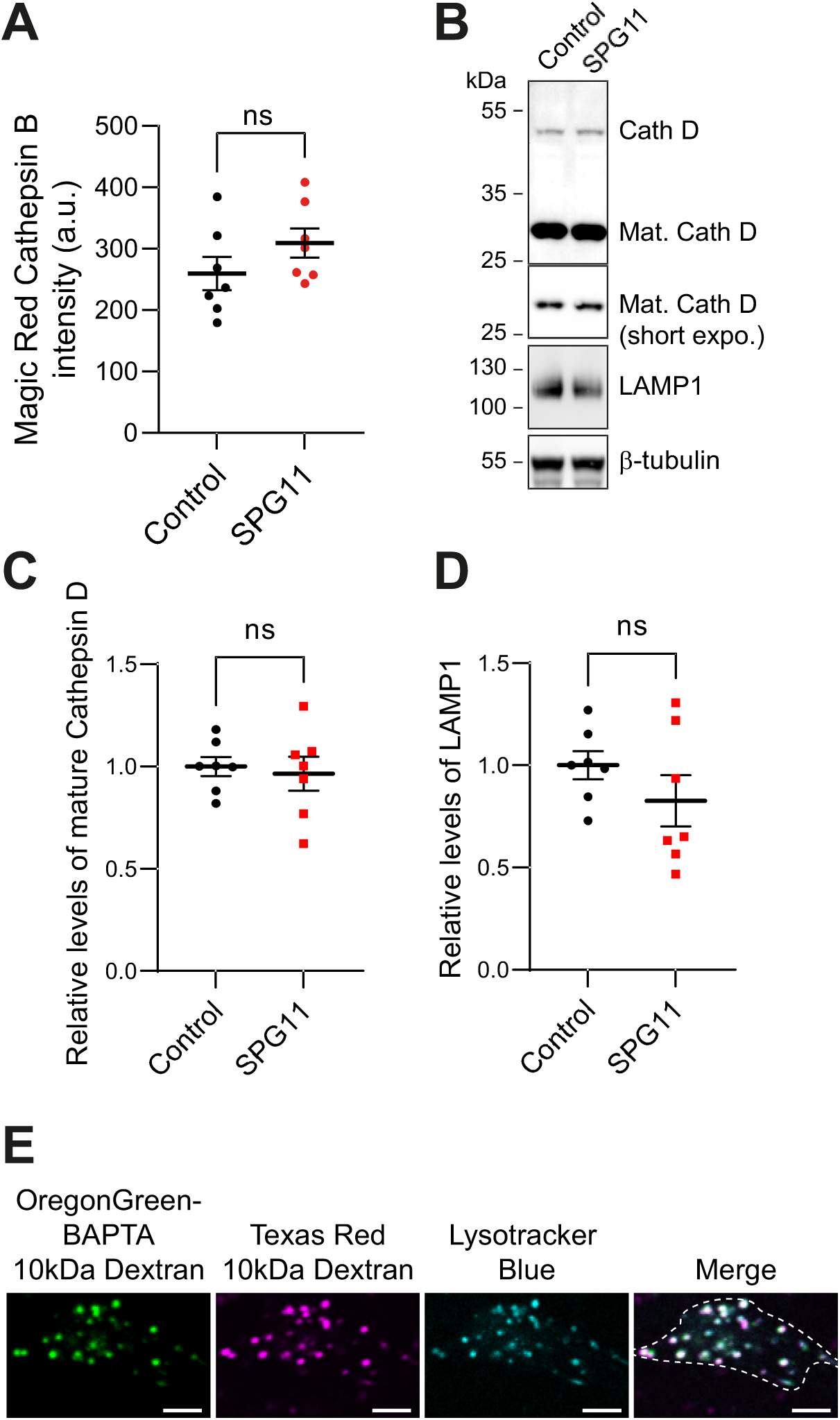
Lysosome catabolic activity is preserved in SPG11 hNPCs. A. Quantification of Magic Red Fluorescent Cathepsin B assay performed on control and SPG11 hNPCs. The graph shows the mean ± SEM. N=7 wells (with >200 cells analyzed per well) of a 24-well plate from 3 independent experiments. Unpaired t-test. ns p=0.19. B. Western blots showing expression levels of the lysosomal proteins LAMP1 and Cathepsin D. For the latter, note the presence of full length, unprocessed Cathepsin D (Cath D), and the cleaved mature Cathepsin D (Mat. Cath D). C. -D. Quantification of the relative levels of the mature form of Cathepsin D and LAMP1, normalized to levels of β-tubulin. The graphs show the mean ± SEM. N=7 independent experiments. Unpaired t-test, ns p>0.24. E. Live confocal image of hNPCs loaded for 16 hours with 10 kDa dextran Texas Red and 10 kDa dextran coupled to the calcium probe Oregon Green 488 BAPTA and stained with Lysotracker blue. Dashed line indicates the cell periphery. Note the colocalization of 10 kDa dextran probes with Lysotracker. Scale bar: 5 µm.

**Supplementary Figure 3.**
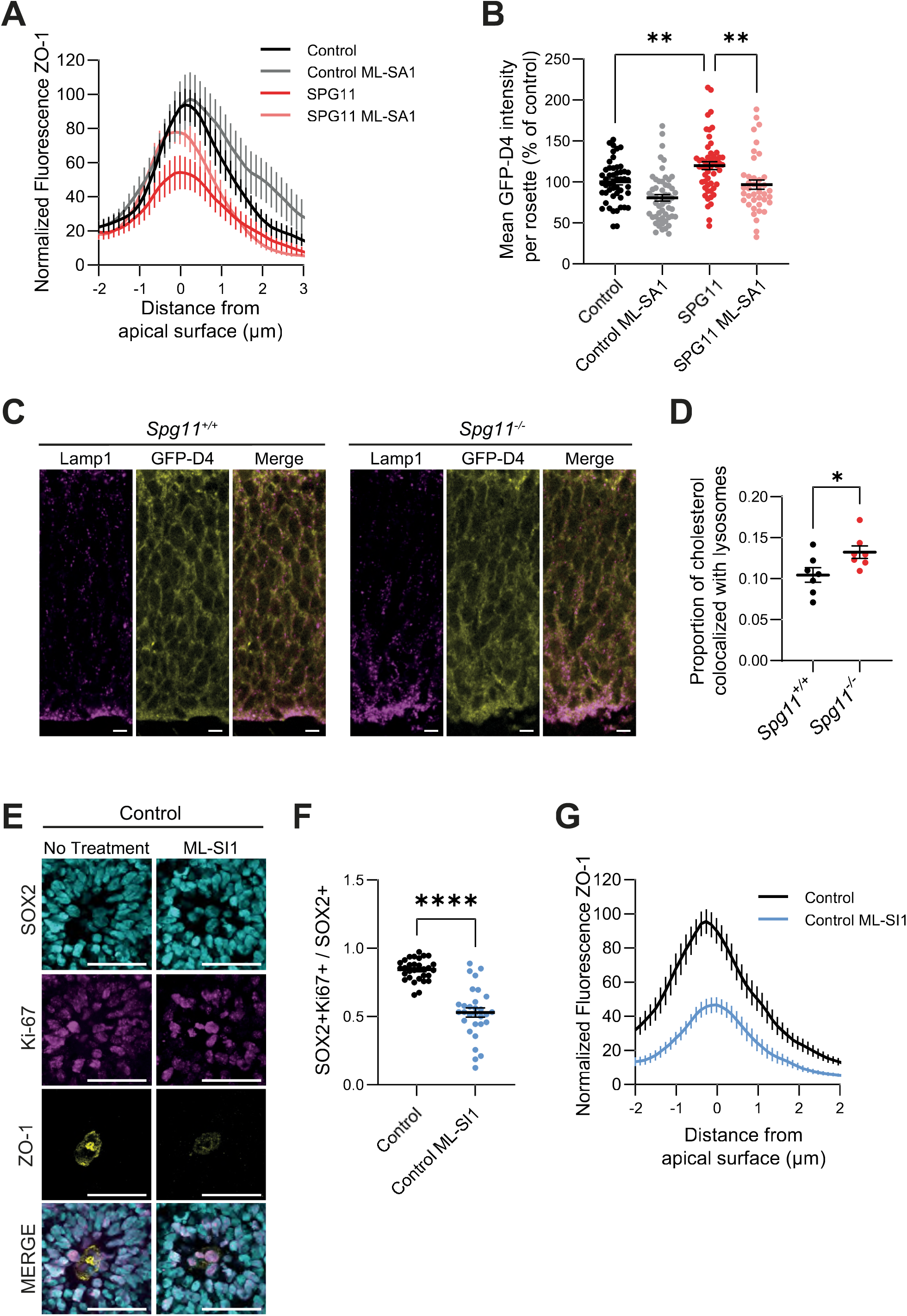
Altering lysosomal calcium in cortical organoids impairs neural progenitor proliferation. A. Quantification of ZO-1 fluorescence along a line scan perpendicular to the apical surface of randomly selected rosettes in control and SPG11 14-day-old cortical organoids, either untreated or treated with ML-SA1, indicating tight junction enrichment in neural progenitors. The graph shows the mean ± SEM. N=30 rosettes from 3 independent experiments. Note the decrease in the intensity of ZO-1 immunostaining at the apical surface of SPG11 rosettes, which is compensated in the presence of ML-SA1 treatment. B. Quantification of the mean intensity of GFP-D4 fluorescence per rosette in control and SPG11 cortical organoids, either untreated or treated with ML-SA1. The graph shows the mean ± SEM. N>40 rosettes from 3 independent experiments. One-way ANOVA followed by Holm-Sidak’s multiple comparisons test, **p<0.01. C. Cortical section of E12.5 old wild-type (*Spg11^+/+^*) or *Spg11* knockout (*Spg11^-/-^*) mice were immunostained with antibody against the lysosomal marker Lamp1, and stained with the GFP-D4 probe to detect cholesterol. Scale bar: 20µm. D. Quantification of the proportion of GFP-D4 staining that is colocalized with lysosomes (Lamp1 immunostaining) in the cortex of E12.5 of wild-type (*Spg11^+/+^*) or *Spg11* knockout (*Spg11^-/-^*) mice. The graph shows the mean ± SEM. N=7 independent animals. Mann-Whitney test. * p<0.05. E. Rosettes of control 14-day-old cortical organoids, either untreated or treated with the TRPML1 antagonist ML-SI1 were immunostained with antibodies against SOX2 for neural progenitors, Ki67 for proliferating cells and ZO-1 for tight junctions. Scale bar: 50 µm. F. Quantification of the proportion of apical SOX2-positive cells that were also positive for the Ki67 marker in control organoids that were untreated or treated with ML-SI1. The graph shows the mean ± SEM. N=30 rosettes from 3 independent experiments. Unpaired t-test, ****p<0.0001. |G. Quantification of ZO-1 fluorescence along a line scan perpendicular to the apical surface of randomly selected rosettes in control 14-day-old cortical organoids, either untreated or treated with ML-SI1. The graph shows the mean ± SEM. N=30 rosettes from 3 independent experiments.

**Supplementary Figure 4.**
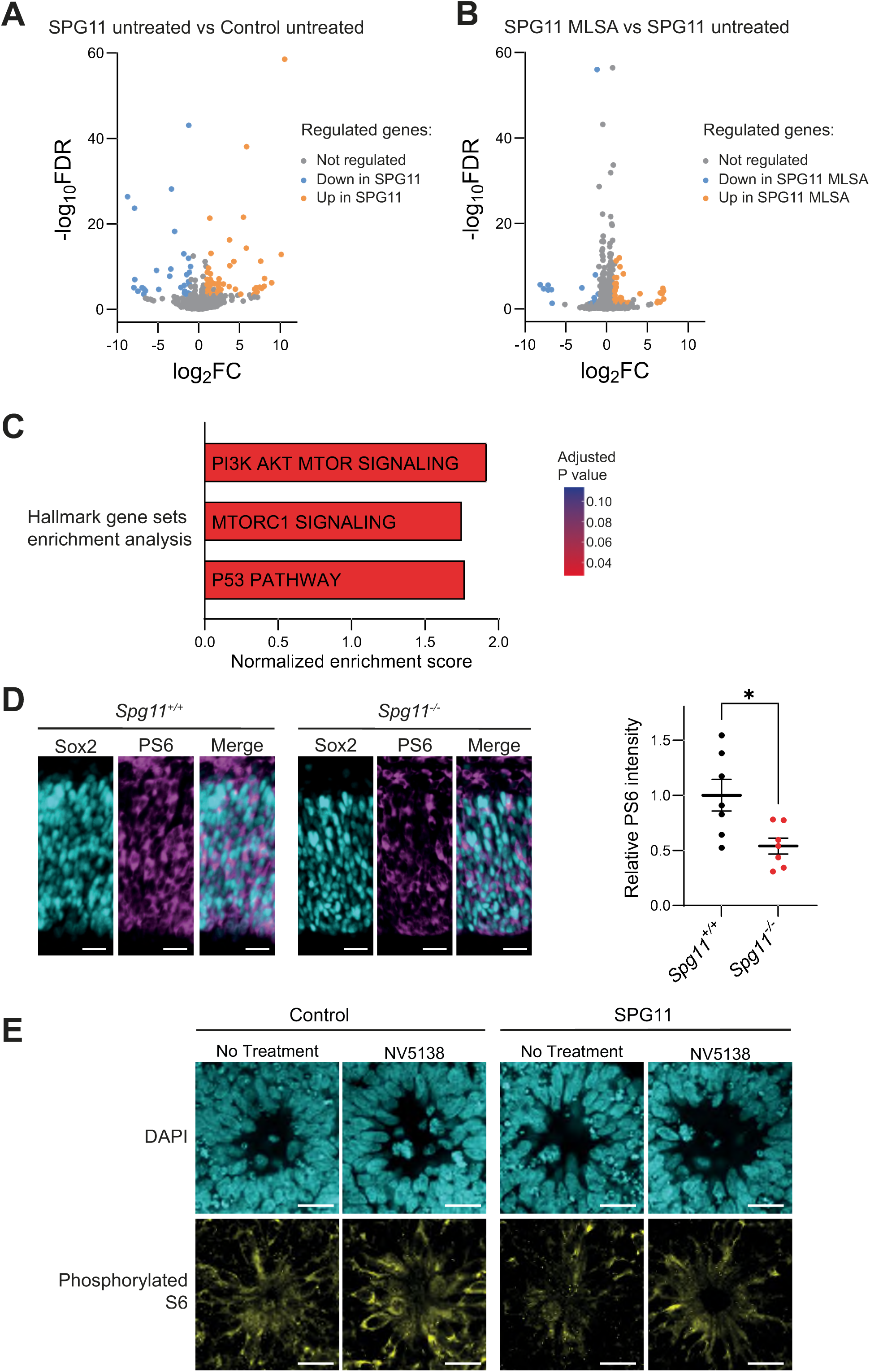
ML-SA1 treatment modulates transcription in SPG11 cortical organoids. A. Volcano plot depicting differential gene expression of transcriptomes between SPG11 and control 14-day-old cortical organoids (N=4). Genes significantly upregulated in SPG11 organoids are orange, while significantly downregulated genes are blue. EdgeR analysis using log2FC > 1, FDR < 0.05. Covariate corrections performed to account for experiments launched separately. B. Volcano plots depicting differential gene expression of transcriptomes between untreated and ML-SA1 treated SPG11 14-day-old cortical organoids (N=4). Genes significantly upregulated by ML-SA1-treated organoids are orange, while significantly downregulated genes are blue. EdgeR analysis using log2FC > 1, FDR < 0.05. Covariate corrections performed to account for experiments launched separately. C. Summary of geneset enrichment analysis (GSEA) results using the MsigDB Hallmark database showing pathways with an adjusted p-value <0.05. D. Cortex of E12.5 control and *Spg11*-knockout (*Spg11^-/-^*) mouse were immunostained with antibodies against Sox2 and phosphorylated S6 (PS6). Scale bar: 20µm. Quantification the relative intensity of PS6 in Sox2-positve progenitors of cortical sections obtained from embryonic day 12.5 wild-type (*Spg11^+/+^*) or *Spg11* knockout (*Spg11^-/-^*) mice. The graphs show the mean ± SEM. N=8 independent animals. Mann-Whitney test, *p<0.05. E. Rosettes in control and SPG11 14-day-old cortical organoids, either untreated or treated with NV5138, were stained with DAPI and immunostained with antibodies against phosphorylated S6. Scale bar: 20 µm.

**Supplementary Figure 5.**
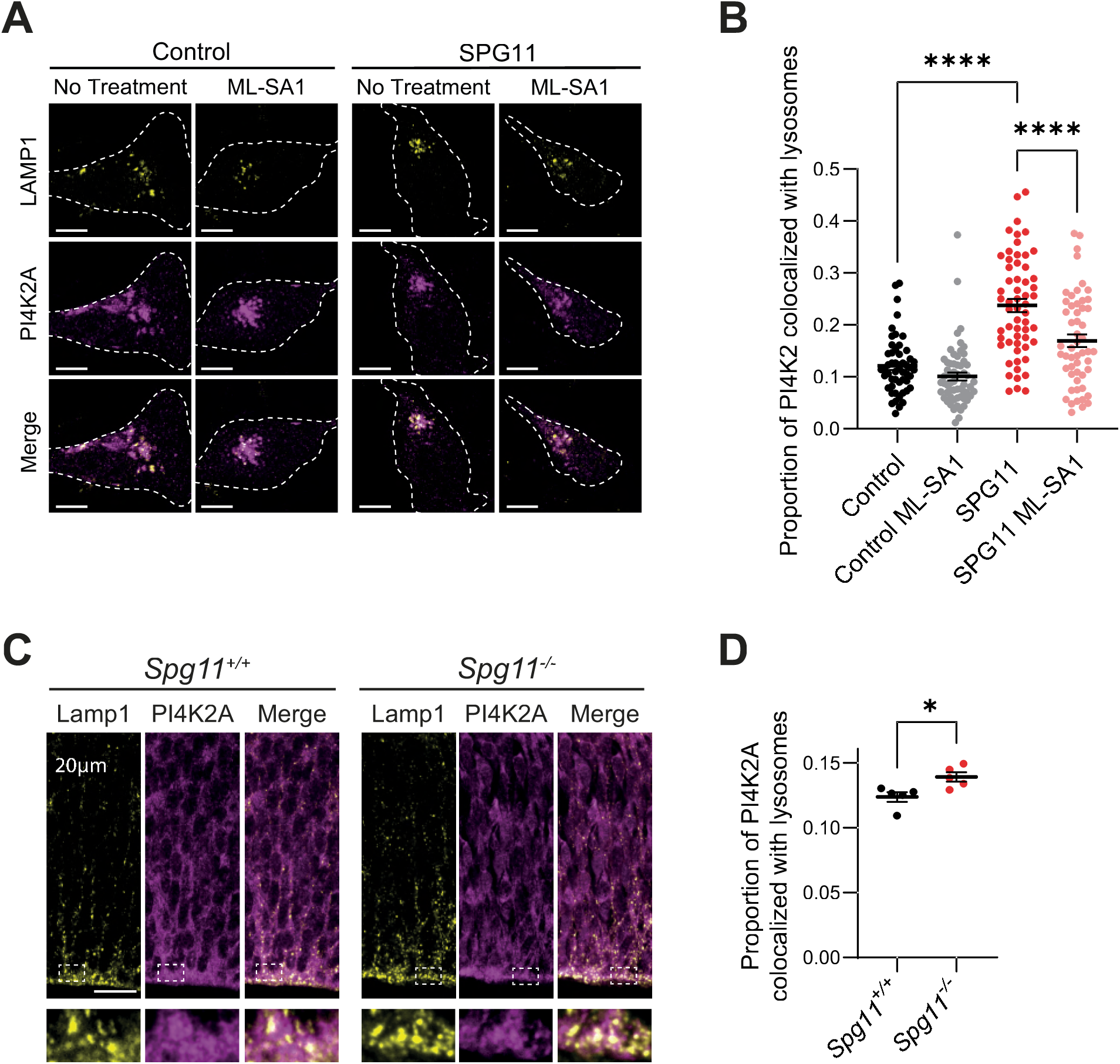
ML-SA1 treatment modulates PI4K2A lysosomal localization in SPG11 hNPCs. A. Immunostaining of control or SPG11 hNPCs, either untreated or treated with ML-SA1, with antibodies raised against PI4K2A and LAMP1 to stain lysosomes. Scale bar: 10 µm. B. Quantification of the proportion of PI4K2A signal colocalized with lysosomes labelled by LAMP1 immunostaining in control or SPG11 hNPCs, either untreated or treated with ML-SA1 for 24 hours. The graph shows the mean ± SEM. N>50 hNPCs analyzed from 3 independent experiments. Ordinary one-way ANOVA followed by Holm-Sidak’s multiple comparisons test, ****p<0.0001. C. Cortical section of E12.5 old wild-type (*Spg11^+/+^*) or *Spg11* knockout (*Spg11^-/-^*) mice were immunostained with antibody against PI4K2A and the lysosomal marker. Scale bar: 20µm. D. Quantification of the proportion of PI4K2A signal colocalized with lysosomes (Lamp1 immunostaining) in the cortex of E12.5 of wild-type (*Spg11^+/+^*) or *Spg11* knockout (*Spg11^-/-^*) mice. The graph shows the mean ± SEM. N=5 independent animals. Mann-Whitney test, * p<0.05.

## Notes

### Competing Interest Statement

The authors have declared no competing interest.

### Summary of Updates

We added data obtained in a Spg11 knockout mouse model (Figure 1, Suppl. Fig. 1) to support the findings in organoid models.

